# Dual functions of Discoidin domain receptor coordinate cell-matrix adhesion and collective polarity in migratory cardiopharyngeal progenitors

**DOI:** 10.1101/154880

**Authors:** Yelena Y. Bernadskaya, Saahil Brahmbhatt, Stephanie E. Gline, Wei Wang, Lionel Christiaen

## Abstract

Integrated analyses of regulated effector genes, cellular processes, and extrinsic signals are required to understand how transcriptional networks coordinate fate specification and cell behavior during embryogenesis. Migratory pairs of cardiac progenitors in the tunicate *Ciona* provide the simplest model of collective migration in chordate embryos. Ciona cardiopharyngeal progenitors (aka trunk ventral cells, TVCs) polarize as leader and trailer cells, and migrate between the ventral epidermis and trunk endoderm, which influences collective polarity. Using functional perturbations and quantitative analyses, we show that the TVC-specific and collagen-binding Discoidin-domain receptor (Ddr) cooperates with Integrin-β1 to promote cell-matrix adhesion to the epidermis. We found that endoderm cells secrete a collagen, Col9-a1, that is deposited in the basal epidermal matrix and activates Ddr at the ventral membrane of migrating TVCs. A functional antagonism between Ddr/Intβ1-mediated cell-matrix adhesion and Vegfr signaling appears to modulate the position of cardiopharyngeal progenitors between the endoderm and epidermis. Finally, we show that Ddr activity promotes leader-trailer-polarized BMP-Smad signaling independently of its role in cell-matrix adhesion. We propose that dual functions of Ddr act downstream of cardiopharyngeal-specific transcriptional inputs to coordinate subcellular processes underlying collective polarity and directed migration.

---

During embryonic development, complex tissue-scale movements emerge from collective behaviors of individual cells. Morphogenesis is tissue-specific, indicating that the gene regulatory networks (GRNs) that control cell identity also determine cell behavior. How GRNs control and coordinate cell fate and behavior has been illustrated using as diverse models as mesoderm invagination and migration in Drosophila and sea urchins, gastrulation in Xenopus, and neural crest cell migration in amniotes^1-8^. Among other essential morphogenetic processes, collective migration is observed in various physiological and pathological conditions such as neural crest cell migration, gastrulation, wound healing, and cancer metastasis^9, 10^. During collective movements, cells can be of the same identity but adopt different leader and follower states, with distinctive morphologies^11^. Collective polarity must be established, maintained and coordinated with the direction of movement, and both polarity and directionality arise from anisotropic exposure to extrinsic cues, such as the free edge of leading cells, gradients of secreted molecules from surrounding tissues, or asymmetric distribution of the adjacent extracellular matrix^12, 13^. Integrative mechanisms must therefore connect regulatory inputs with the production of state-specific cell behavior in response to extrinsic cues.

The cardiac lineage in the tunicate *Ciona* provides the simplest example of collective cell migration^13-16^. In *Ciona*, multipotent cardiopharyngeal progenitors derive from the B7.5 blastomeres in 110-cell embryos, and later produce fate-restricted heart and pharyngeal muscle precursors^17-20^. On either side of tailbud embryos, pairs of cardiopharyngeal progenitor cells (aka trunk ventral cells, TVCs) collectively polarize and migrate between the ventral epidermis and the trunk endoderm, until they stop and produce distinct fate-restricted progenitors ^13, 14, 16, 19^. During migration, the leader TVC extends dynamic protrusions and generates a broad leading edge, while the trailer terminates in a tapered retraction end^13, 16^. Before migration, the surrounding trunk endoderm preferentially contacts the prospective leader cell, and experimental perturbation of secretion in endoderm cells disrupts collective leader-trailer (LT) polarity^13^, suggesting that anisotropic exposure to biochemical cues secreted by the endoderm contributes to collective TVC polarity. Moreover, B7.5-lineage-specific transcriptional inputs from Mesp, FGF/MAPK signaling and Foxf control and coordinate cardiopharyngeal fate specification and TVC migration^15, 16, 18, 21^, and the transcriptome of migratory TVCs have been extensively profiled^16, 22-24^. Therefore, TVC migration provides an attractive model to functionally connect fate-specific transcriptional inputs with cellular effectors and extrinsic cues governing collective polarity and directed migration.

Among regulated cellular effectors possibly shaping cell responses to extrinsic cues, we identified several receptor tyrosine kinases (RTKs)^16^. Other developmental cell behaviors have been shown to involve RTK signaling, including for collective polarization of migratory cell groups in vertebrates and Drosophila^10, 25-30^. We thus selected a small group of candidate RTKs, and analyzed their roles in TVC migration. Among these candidates, the sole Ciona homolog of *Discoidin domain receptor* (*Ddr*) appeared to be upregulated specifically in newborn TVCs, downstream of both FGF/MAPK signaling and Foxf inputs^16^. DDRs are single pass transmembrane receptors that bind extracellular collagen^31, 32^, they can mediate weak adhesion to collagen and regulate cadherins, integrins and ECM interacting proteins to modulate cell-matrix adhesion^33-35^.

Here we describe the functions of Ddr and its interactions with integrin, Vegfr and BMP-Smad signaling in regulating cell-matrix adhesion and collective polarity during TVC migration. We developed methods to quantify TVC morphology and movements, and defined an experimental and analytical framework to study morphogenetic determinants. We found that the endoderm secretes a type IX collagen, Col9-a1, that is deposited onto the basal epidermal matrix and activates Ddr at the ventral surface of the migrating TVCs, thus promoting integrin-based cell-matrix adhesion. We dissected a signaling antagonism between Ddr/Integrin-mediated cell-matrix adhesion and Vegfr signaling. Finally, we show that Ddr promotes polarized BMP-Smad signaling, thus contributing to establishing distinct leader and trailer states. The latter function appears independent from integrin-mediated cell-matrix adhesion, suggesting that dual functions of Ddr coordinate collective polarity and cell-matrix adhesion during TVC migration.

## RESULTS

### Cardiopharyngeal progenitors express Receptor Tyrosine Kinases

Transcriptome profiling identified candidate signaling molecules potentially involved in guiding the movement of Ciona cardiopharyngeal progenitors, the TVCs. We used whole mount *in situ* hybridization to confirm and better characterize the expression of four candidate RTK-coding genes, *Discoidin domain receptor* (*Ddr*), *Vascular endothelium growth factor receptor* (*Vegfr*)*, Fibroblast growth factor receptor* (*Fgfr*), and epidermal growth factor receptor (*Egfr)*. We did not detect *Egfr* transcripts prior to or during TVC migration, neither did it show signs of being functional in overexpression of truncated dominant negative forms (Figure S1, Movie 5), we thus excluded Egfr from further analysis (Figure S1). *Ddr*, *Vegfr*, and *Fgfr* transcripts were detected in migrating TVCs (Figures 1B, S1). *Vegfr* and *Fgfr* were expressed in B7.5 lineage founder cells, while newborn TVCs upregulated *Ddr* before the onset of collective migration (Figure S1).

The transcription factor Foxf was proposed to act as a key transcriptional regulator of TVC migration^15^, and transcription profiling identified candidate target genes, including *Ddr*^16^. Using B7.5-lineage-specific CRISPR/Cas9-mediated mutagenesis with *Foxf* targeting single guide RNAs (sgRNAs;^36^), we confirmed that loss of Foxf function inhibited *Ddr* expression in the TVCs (Figure 1B,C). By contrast, Foxf function appeared dispensable for *Vegfr* or *Fgfr* expression, a finding also consistent with previous microarray analyses^16^. Parallel studies indicated that Fgfr is primarily required for MAPK-dependent transcriptional regulation in migrating TVCs and beyond^37^. Taken together, these data led us to focus on Ddr and Vegfr as candidate cell migration effectors, which we sought to further characterize.

**Figure 1.**
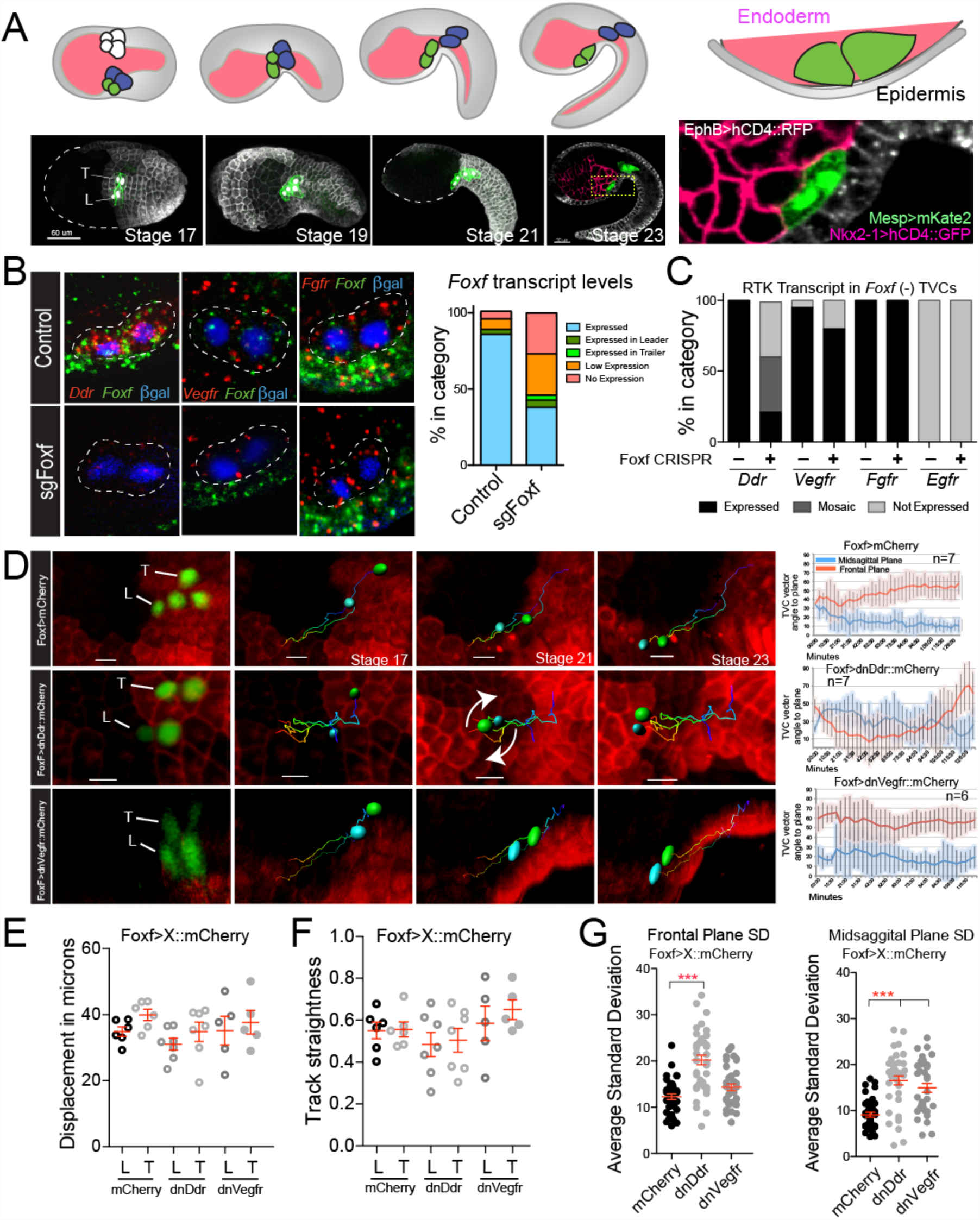
Ddr and Vegfr regulate trunk ventral cell (TVC) migration. **A**. Schematic of TVC migration between the endoderm and epidermis. Cousin TVCs in green, ATMs are blue, endoderm in pink, and epidermis in grey. Micrographs of TVC migration and tissue contact. Epidermal cells are marked with *EfnB>hCD4:: mCherry*, B7.5 lineage marked with *Mesp>hCD4:: GFP* and nuclear *Mesp>H2B:: mCherry* for stages 17-19, markers used for 3-channel acquisition are given on the magnified image. Close-up shows the positioning of the TVCs between the endoderm and epidermis. **B**. Fluorescent *in situ* hybridization (FISH) carried out in migrating TVCs at stage 23 in control and CRISPR targeting the *Foxf* locus for mutagenesis. All micrographs are oriented with the leader TVC to the left. TVC pair is outlined with dotted line and nuclei are marked with Mesp>LacZ and stained for β-galactose. Quantification of *Foxf* knockout efficiency is shown to the right. **C**. RTK expression in the proportion of embryos that lose *Foxf* expression. Guide RNAs targeting the *Ebf* locus for mutagenesis are uses as controls as *Ebf* is not required for TVC migration and does not affect expression of TVC-specific genes. **D**. TVC positional tracking. TVC (and ATM) nuclei are marked with *Mesp>H2B:: GFP* and epidermis with *EfnB>hCD4:: mCherry*. Tracks of migrating TVCs are shown in each panel, cyan sphere = leader, green sphere = trailer. Arrows show the tumbling of the TVCs as leader and trailer switch positions. L = leader, T= Trailer. Average position angle to sagittal and frontal planes are shown with standard deviation. **E**. Average displacement of leader and trailer TVCs during migration. **F.** Track straightness of leader and trailer TVCs. **G**. Average standard deviation at the sagittal and frontal planes. *** = p<0.0005.

### Ddr and Vegfr functions are required for proper TVC migration

Wild type TVC migration is a highly stereotypical process. Soon after birth, pairs of cousin TVCs collectively polarize to assume leader and trailer positions, potentially through differential contacts with the mesenchyme and the trunk endoderm^13^, and these relative positions are maintained throughout migration. The cells move anteriorly, between the ventral epidermis and trunk endoderm^13, 16^, until they reach a final position adjacent to the midline, where they divide asymmetrically to generate the first and second heart precursors, and atrial siphon muscle precursors^19, 20^. To better characterize TVC migration in control and experimental conditions, we developed quantitative parameters describing cell movements in live embryos. We used an epidermal transgene, *EfnB>hCD4::mCherry*^13^, and a B7.5-lineage-specific nuclear marker, *Mesp>H2B::GFP*, to label live embryos and migrating cardiopharyngeal progenitors and imaged them using time lapse confocal microscopy for the duration of TVC migration (Figure 1D, movies S1-S5). To quantify TVC movements in four dimensions, we used morphological landmarks in the epidermis to define sagittal and frontal planes. We used GFP+ TVC nuclei to define the leader-trailer (LT) axis, and calculated the angles it formed with sagittal and frontal planes at each time point (Figures 1D, S2A, Methods). We used cell division times to align temporal axes and combine time-lapse recordings from multiple embryos (n=7). In this way, we found that the TVC migration path is highly constrained in control embryos, with LT angles with frontal and sagittal planes changing within defined ranges as cells migrate (Figures 1D,G, S2).

To study the functions of selected RTKs during TVC migration, we generated passive dominant negative versions of Ddr and Vegfr (henceforth referred to as dnDdr and dnVegfr), by deleting the cytoplasmic kinase domain of each receptor. We expressed these constructs in the newborn TVCs using a defined *Foxf* enhancer^15^, and assayed migration quantitatively. TVCs expressing *Foxf*-driven dnDdr retained their ability to initiate migration, but cells tumbled, failing to maintain relative leader/trailer positions and followed more variable migration paths (Figure 1D-G, Movie S2). DnDdr misexpression also caused a decrease in total displacement of TVCs and in path straightness (Figure 1E,F) consistent with altered directionality (Supplemental Fig. 2). These observations indicate that proper function of the Foxf target Ddr is required for maintenance of collective polarity and directional migration of the TVCs.

TVCs expressing *Foxf*-driven dnVegfr were also able to initiate migration, and their total displacement was comparable to control cells (Figure 1G). However, dnVegfr expression increased track straightness, as if extrinsic constraints canalized migration more efficiently than in control embryos^13^ (Figure 1F). To quantify the variability of cell behaviors, we calculated the standard deviation of the angles between LT axes and the sagittal and frontal planes at each time point during migration. As expected, dnDdr increased the standard deviation of the LT angle relative to both the sagittal and the frontal plane, reflecting the variable positions of tumbling TVCs. By contrast, dnVegfr only increased the standard deviation of the LT angle relative to the sagittal plane, which is consistent with cells remaining constrained in at least one dimension (Figure 1G).

### Ddr promotes integrin-based cell-matrix adhesion to the epidermis

We first focused on understanding how Ddr regulates TVC polarity and migration, and analyzed changes in cell morphology and contacts with surrounding tissues^13^. We used defined transgenes to label TVCs’ nuclei and membranes, as well as surrounding tissues, and used confocal imaging and computational segmentation of individual cells to quantify morphometric parameters, including cell sphericity and the percentage of total cell surface contacting the underlying epidermis. By revealing quantitative differences between leader and trailer cells, these measurements allowed us to characterize collective polarity in control and experimental conditions.

Sphericity is calculated as the ratio between the surface area of a sphere of the same volume as the object of interest and the surface area of that object. The maximum sphericity would thus be 1 if the cell was a perfect sphere, and it decreases in migratory cells that protrude and flatten on a 2D substrate. Our analysis finds that the leader TVC is less spherical than the trailer, which is consistent with its greater protrusive activity16. Approximately 20% of the leader cell surface engages in the cell-cell junction with the trailer (Figure S3). At the same time, 45-50% of the leader surface contacts the underlying epidermis compared to 40-45% for the trailer (Figure 2A,C). Expression of dnDdr significantly increased the sphericity of the leader, abolishing leader-trailer differences (Figure 2A,B). Concurrently, dnDdr significantly reduced the surface of TVC contact with the underlying epidermis (Figure 2A,C). dnDdr-expressing TVCs contacted the epidermis with about 30% of their surface on average with 35% of the cells completely detaching from the epidermis. Complementary loss-of-function approaches validated the specificity of the dnDdr construct, as lineage-specific RNAi using short hairpin microRNA (shmiR) constructs synergized with suboptimal doses of dnDdr to cause a penetrant “de-adhesion” phenotype (Figure S4). These data suggest that Ddr is required for the TVCs to maintain contact with the epidermis during migration.

**Figure 2.**
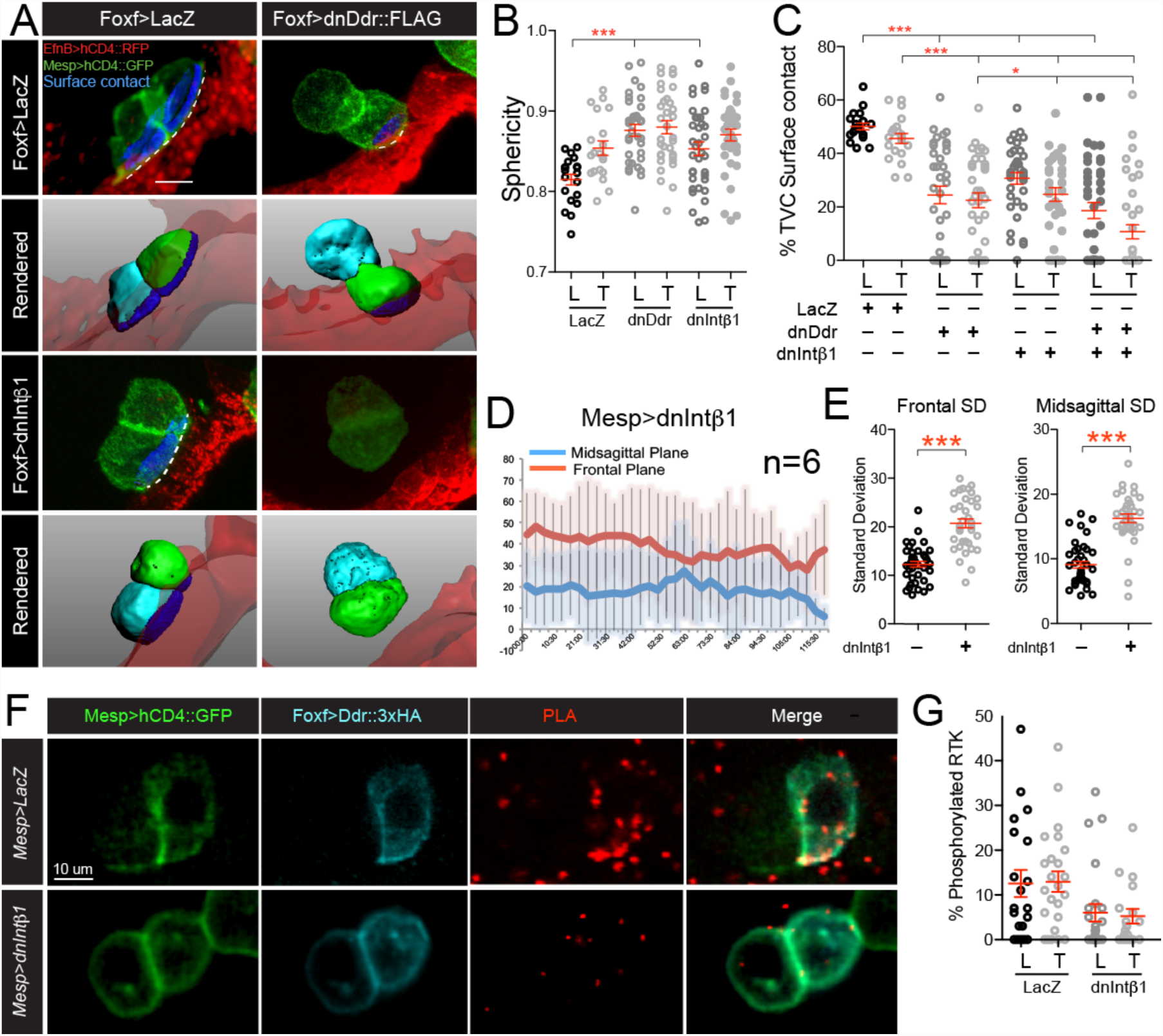
Integrin-β1 and Ddr promote TVC adhesion to the ventral epidermis. **A**. Micrographs of surface contacts (blue) made by migrating TVCs with the underlying epidermis. On the upper panels, the length of the surface contact is indicated with a dotted line. TVCs are marked and segmented based on the B7.5 linage-specific expression of *Mesp>hCD4::GFP* and epidermal surface is visualized with *EfnB>hCD4:: tagRFP*. Lower panels show rendered surfaces, leader is shown in cyan, trailer in green, epidermal surface in transparent red. For segmentation and derivation of surface contacts see Materials and Methods. **B**. Morphology of TVC under perturbation conditions measured as sphericity of Leader/Trailer cells during migration. **C**. Quantitation of percent TVC surface in contact with the epidermis. All transgenes are expressed from the Foxf TVC-specific enhancer. Results shown are pooled from 3 biological replicate experiments. **D**. Quantitation of migration angles of *Mesp>dnIntβ1*- expressing TVCs, averages at each time point and standard deviation shown. **E**. Average standard deviation of the angle between the LT axis and the frontal and sagittal planes in *Mesp>dnIntβ1*-expressing TVCs. **F**. Proximity ligation assay (PLA) of phosphorylated full-length Ddr under control and dnIntβ1 conditions. **G**. Quantitation of phosphorylated Ddr compared to the total amount of Ddr. All images are oriented with leader TVC to the left. L=leader, T=Trailer. * = p<0.05, ** = p<0.005, *** = p<0.0005.

Phenotypes produced by dnDdr suggested that cells failed to maintain adhesion to the extracellular matrix lining the ventral epidermis. For instance, Ddr homologs can mediate weak adhesion to collagen, and increase cells’ adhesion to collagens through integrins^34, 38^. We thus sought to test whether Ddr contributes to cell-matrix adhesion, and identify its possible integrin partners in the migrating TVCs. Using whole mount fluorescent *in situ* hybridization assays and TVC-specific transcriptome profiling data^16^, we detected expression of Integrin-β1 (Intβ1), among several other candidates (Supplementary Figure 5). We generated a dominant negative form of Intβ1 (dnIntβ1) by truncating the C-terminal cytoplasmic domain^39^, and assayed cell sphericity and the ability of dnIntβ1-expressing TVCs to contact the epidermis (Figure 2A-C). Altering Intβ1 function increased cells’ sphericity to levels comparable to those observed with dnDdr (Figure 2B). Furthermore, dnIntβ1-expressing TVCs contacted the epidermis with only 20 to 30% of their surface. Compared to control cells, this reduction is similar to that observed with dnDdr. Finally, dnIntβ1 also increased the variability of TVC position during migration, similar to dnDdr (Figure 2D). Taken together, these data indicate that altering Integrin-β1 and Ddr functions cause similar phenotypes characterized by a loss of adhesion to the ventral epidermis, cell rounding and unstable collective polarity. We thus conclude that Ddr, like Integrin-β1, functions primarily to promote cell-matrix adhesion at the contact with the ventral epidermis, and this adhesion stabilizes collective polarity, permitting directed migration.

Since both Ddr and Intβ1 can function as collagen receptors ^31, 32^, we sought to test whether they interact to regulate cell-matrix adhesion. We quantified TVCs’ contacts with the epidermis following dnDdr and/or dnIntβ1 expression. Co-expression of suboptimal doses of dnDdr and dnIntβ1 aggravated the detachment phenotype, but to a lesser extent than expected for additive effects (Figure 2A,C). This suggests that Ddr and Intβ1 function primarily in overlapping pathways to regulate TVC/epidermis adhesion. To further characterize the functional relationships between the two collagen-receptors, we asked whether Intβ1 is required for Ddr activation. To assay localization and activation of full-length Ddr, we used the minimal *Foxf* TVC enhancer and expressed a 3xHA-tagged version of Ddr at minimal detectable levels, to avoid non-specific localization and over-expression phenotypes. We then performed immunohistochemisty (IHC) combined with a proximity ligation assay (PLA) using anti-HA and anti-phospho-Tyrosine antibodies to visualize the phosphorylated form of Ddr^38, 40^. Full-length Ddr::3xHA proteins were localized to both intracellular vesicles and the ventral plasma membrane (Figures 2E). However, activated Ddr preferentially localized to the ventral TVC surface, which contacts the epidermis (Figure 2E). Quantitative analysis showed that, at any given time, 12 to 20% of tagged Ddr proteins were phosphorylated, suggesting that there is a large pool of inactive Ddr proteins within the cells.

To test whether Ddr activation on the ventral/epidermal side of migrating TVCs requires integrin-based cell-matrix adhesion, we repeated the PLA assays in embryos expressing the B7.5 lineage-specific *Mesp>dnIntβ1* transgene. We found that dnIntβ1 misexpression reduced Ddr phosphorylation levels to approximately 6%, and the enrichment of activated Ddr on the ventral cell surface of the TVCs was lost (Figure 2E,F), indicating that Intβ1 activity is required to localize and activate Ddr on the ventral/epidermal side of migrating TVCs. Taken together, these observations reinforce the notion that Ddr and Intβ1 interact to promote TVC adhesion to the extracellular matrix on the epidermal side of the cells.

### The endoderm contributes to cell-matrix adhesion and Ddr activation

Previous work showed that expressing a dominant negative form of the small GTPase Sar1 (dnSar1) with the endoderm-specific *Nkx2-1* enhancer inhibited ER-to-Golgi transport and secretion from the endoderm, and caused a tumbling phenotype reminiscent of that observed with dnDdr and dnIntβ1^13^. To determine if the *Nkx2-1>dnSar1* construct caused the TVCs to detach from the epidermal matrix during migration, we quantified contacts between the TVCs and the epidermis. As was observed with TVC-specific dnDdr/dnIntβ1 misexpression, *Nkx2-1>dnSar1*-mediated inhibition of secretion in the endoderm caused a significant reduction of the TVC surface contact with the epidermis (Figure 3B), suggesting secretion from the endoderm enables cell-matrix adhesion between the migrating TVCs and the ventral epidermis.

**Figure 3.**
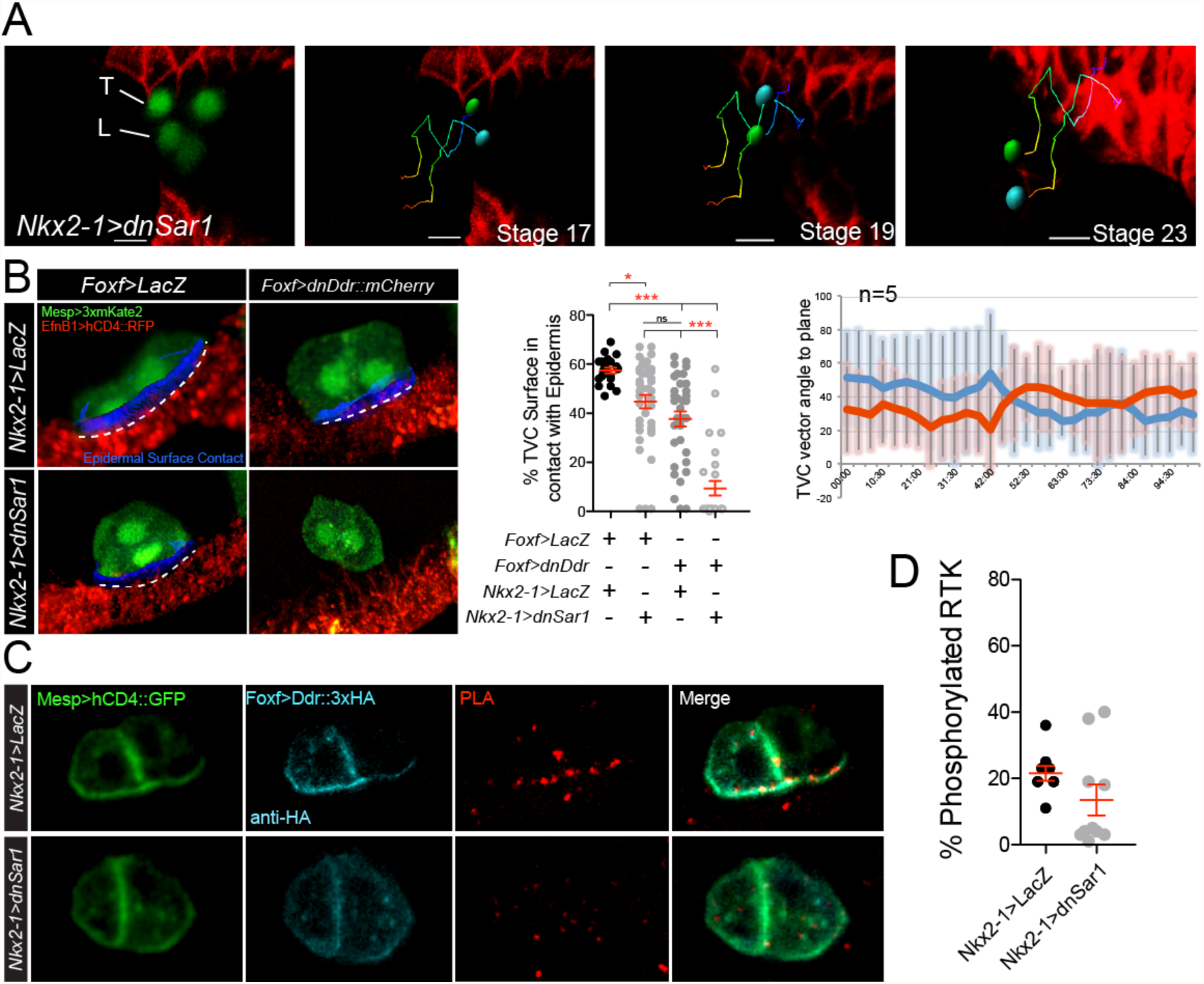
An endodermal cue potentiates TVC adhesion and Ddr activation. **A**. Positional tracking of TVC migration in embryos expressing dnSar1 (*Nkx2-1>dnSar1*) to block secretion in the endoderm. Tracks are time coded from early (blue) to late (red). Leader = cyan, trailer = green. Quantitation of migration angles to frontal and sagittal planes is shown as averages at each time point and standard deviation from average. **B**. Percent of TVC pair surface in contact with epidermis. The B7.5 lineage is marked with *Mesp>3xmKate2*, epidermis is marked with *EfnB>hCD4::tagRFP*. Surface of contact is shown in blue and marked with a dashed line. Scatter plot shows average surface contact under each condition. **C**. PLA of *Foxf>Ddr::3xHA* in control (*Nkx2-1>LacZ*) and *Nkx2-1>dnSar1* conditions. **D**. Quantitation of percent phosphorylated Ddr. All images are oriented with leader TVC to the left. L=leader, T=Trailer. * = p<0.05, ** = p<0.005, *** = p<0.0005.

To test whether endodermal cues potentiate Ddr function in the TVCs, we combined secretion inhibition in the endoderm with TVC-specific dnDdr misexpression and quantified contacts between the TVCs and the epidermis. We observed a significant enhancement of the TVC detachment phenotype by combining *Nkx2-1>dnSar1* and *Foxf>dnDdr* compared to the sum of either perturbation alone (Figure 3B,C), suggesting that the secreted endodermal cue and Ddr function in partially redundant pathways. To further probe the functional interaction between the endoderm and Ddr, we tested if secretion from the endoderm is required for Ddr activation in migrating TVCs. PLA analysis revealed that loss of endodermal secretion decreased the fraction of phosphorylated Ddr on the ventral/epidermal side of migrating cells (Figure 3C,D), suggesting a functional relationship between (a) secreted endodermal cue(s) and the collagen receptor Ddr.

### Endodermal *Col9-a1* is required for Ddr activation and cell-matrix adhesion at the TVC/epidermis interface

As Ddr is a collagen receptor, we hypothesized that that potential endodermal secreted cues could include collagens. *Col9-a1* expression was reported in the endoderm of developing *Ciona* embryos where it contributed to the development of the intestine^41^. We confirmed that *Col9-a1* is expressed in the endoderm adjacent to the ventral epidermis prior to the onset of TVC migration (Figure 4A), thus potentially acting as a source of extracellular collagen for subsequent TVC migration. Other *Col9-a1* isoforms are also expressed strongly in the notochord and the endodermal strand that runs the length of the tail (Figure 4A).

**Figure 4.**
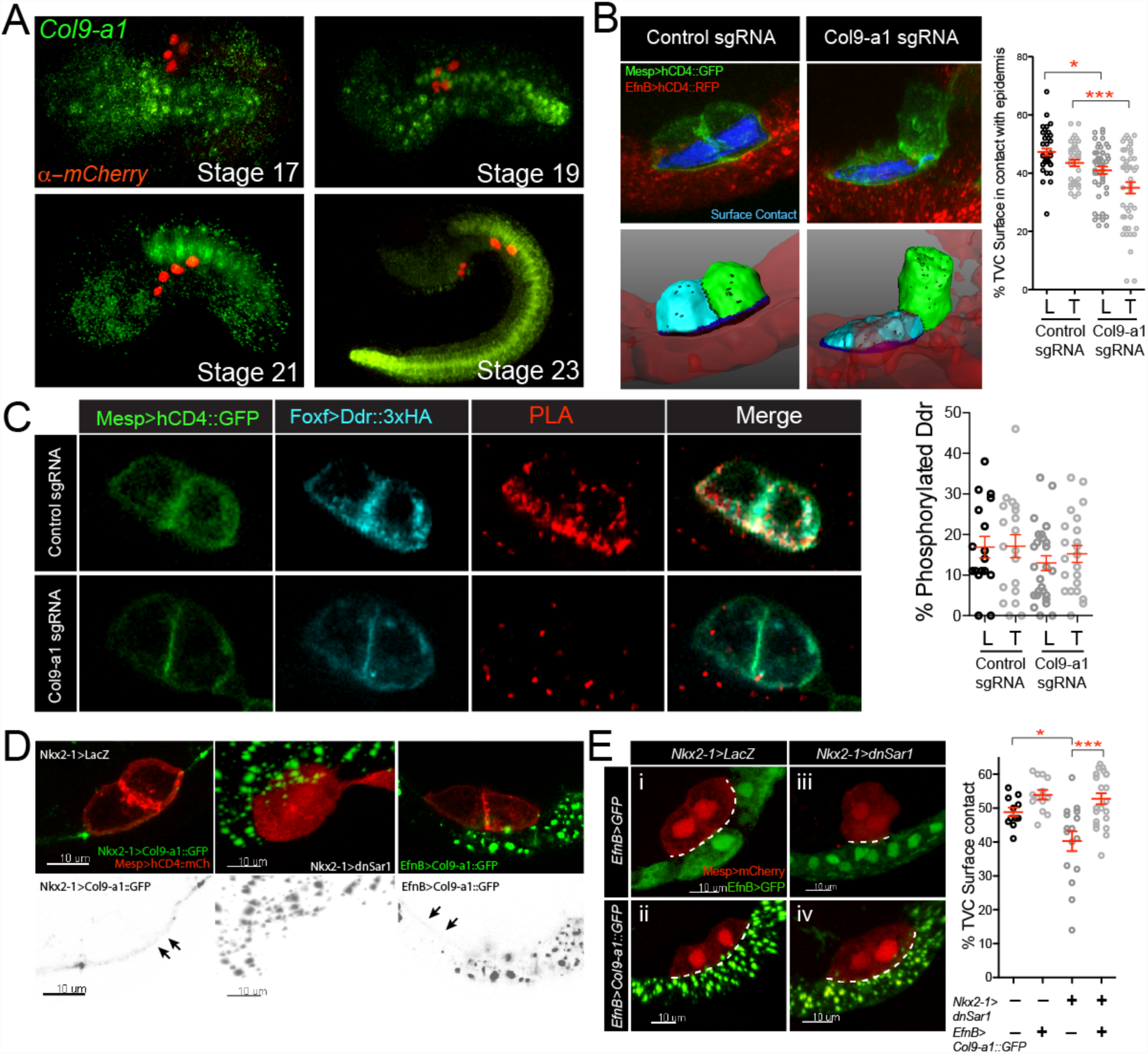
Col9-a1 is the endodermal cue that promotes TVC adhesion and Ddr activation. **A**. Endogenous expression of Col9-a1 in the developing embryo. Micrographs are oriented with anterior to the left, an *in situ* RNA probe is used to visualize *Col9-a1* transcripts. B7.5 lineage is visualized using *Mesp>H2B:: mCherry*. **B**. CRISPR mutagenesis of the *Col9-a1* locus in the endoderm using a *FoxD*-driven Cas9. Blue surfaces are sites of contact. Quantitation of TVC surface area in contact with epidermis adhesion is shown on the right. **C**. PLA to detect phosphorylated full-length Ddr in control (Ebf CRISPR) and CRISPR conditions targeting the *Col9-a1* locus for mutagenesis. **D**. Secretion and localization of a Col9A::GFP fusion expressed in the endoderm (*Nkx2-1*) or in the epidermis (*EfnB*). TVC membrane is visualized using *Mesp>hCD4::mCherry*. Middle panel shows secretion inhibition in the endoderm using dnSar1. Gray scale panels show the corresponding inverted GFP channel. Arrows point to the accumulation of Col9-a1 in the extracellular matrix. **E**. Rescue of *Nkx2-1>dnSar1* TVC adhesion defects by expressing Col9-a1::GFP from the epidermais. Control epidermis is marked with *EfnB>GFP*. Note that in the lower panels EfnB1>GFP is expressed but is less bright in comparison to the Col9-a1::GFP trafficking vesicles. Dotted line outlines the surface of contact. Quantitation of TVC adhesion is shown to the right of the micrographs. All images are oriented with leader TVC to the left. L=leader, T=Trailer. * = p<0.05, ** = p<0.005, *** = p<0.0005.

To test whether Col9-a1 is the endoderm-derived extracellular cue that activates Ddr and promotes cell-matrix adhesion to the ventral epidermis, we used CRISPR/Cas9 to mutagenize *Col9-a1* in the endoderm by expressing Cas9 using an early vegetal enhancer from the gene *FoxD*^42-44^. Two single guide RNAs (sgRNAs) targeting Col9-a1 exons were used to generate large deletions in the gene, and we assayed contacts between the TVCs and ventral epidermis. Col9-a1 gene inactivation in vegetal endoderm progenitors produced a detachment phenotype that mimicked the dnDdr and dnSar1 phenotypes, indicating that collagen Col9-a1 secretion from the endoderm is required for cell-matrix adhesion on the ventral epidermal side of migrating TVCs (Figure 4B).

Since loss of Col9-a1 function phenocopied dnDdr misexpression, and Ddr presumably bind collagens, we sought to test whether endodermal Col9-a1 is required to activate Ddr in the TVCs. We used CRISPR/Cas9 to mutagenize *Col9-a1* in the endoderm and quantified Ddr activation by proximity ligation assay as described above. Loss of Col9-a1 in the endoderm reduced Ddr activation and virtually abolished its localization at ventral membrane contacting the epidermis (Figure 4C), as previously shown upon inhibition of endodermal secretion. Therefore, Col9-a1 is likely the collagen ligand responsible for Ddr activation in the TVCs.

As TVCs migrate into the trunk, they intercalate between the endoderm, which deforms and envelops their dorsal surface, and the epidermis, which remains ventral and seemingly provides the substrate for TVC migration^13, 16^. Since Col9-a1 appears to originate from the endoderm, but activate Ddr on the ventral side where TVCs contact the epidermis, we sought to visualize Col9-a1 localization in the extracellular matrix. We generated a Col9-a1::GFP fusion, which we expressed in the endoderm using the *Nkx2-1* enhancer (*Nkx2-1>Col9-a1-1::GFP*). Col9-a1::GFP was readily secreted from the endoderm and accumulated in the extracellular matrix (Figure 4C). We found that, although the endoderm was the only source of Col9-a1::GFP in these embryos, the GFP signal was detectable in-between the migrating TVCs and the ventral epidermis, whereas the TVC/endoderm interface was devoid of Col9-a1::GFP. We verified that all Col9-a1::GFP proteins accumulated in large vesicles inside endoderm cells upon misexpression of dnSar1 (Figure 4D). This result is consistent with the possibility that endoderm-specific expression of dnSar1 blocks Col9-a1 secretion, which could explain the effects of *Nkx2-1>dnSar1* on Ddr activation and TVC-matrix adhesion.

We attempted to rescue the *Nkx2-1>dnSar1*-induced phenotypes by providing Col9-a1::GFP from an alternative source. Expressing Col9-a1::GFP from the epidermis using an *EfnB* driver^13^ restored the extracellular GFP signal in the matrix lining the ventral epidermis (Figure 4D, gray scale panel). We then quantified contacts between TVCs and the ventral epidermis following misexpression of dnSar1 in the endoderm, together with *EfnB>Col9-a1::GFP* or *EfnB>GFP* as control. Remarkably, resupplying Col9-a1 to the epidermal ECM was sufficient to rescue the TVC adhesion phenotype induced by *Nkx2-1>dnSar1* expression (Figure 4E). This demonstrates that a primary role for the endoderm in TVC migration is to deposit Col9-a1 onto the ventral epidermal ECM, most likely prior to the migration of the TVCs, which then rely on this cue to activate Ddr and adhere to their epidermal substrate.

### An antagonism between Vegfr signaling and Col9-a1/Ddr/Intβ1 functions modulate TVC adhesion to the epidermal matrix

In principle, excessive cell-matrix adhesion would cause TVCs to flatten onto to the ventral epidermis. We occasionally observed this phenotype following TVC-specific misexpression of dnVegfr, although this effect was subtle and did not significantly change cell sphericity (Figures 5A-C). Since dnVegfr expression altered TVC migration (Figure 1), and VEGF receptors and integrin complexes interact functionally in various contexts^45-47^, we sought to test whether Vegfr signaling modulates Ddr/Intβ1-mediated TVC-matrix adhesion. Co-expression of dnVegfr partially suppressed the de-adhesion phenotypes produced by either dnDdr or dnIntβ1 in the TVCs (Figure 5A,B), suggesting that a steady-state antagonism between baseline levels of Vegfr and Ddr/Intβ1 signaling control TVC adhesion to the epidermal matrix.

**Figure 5.**
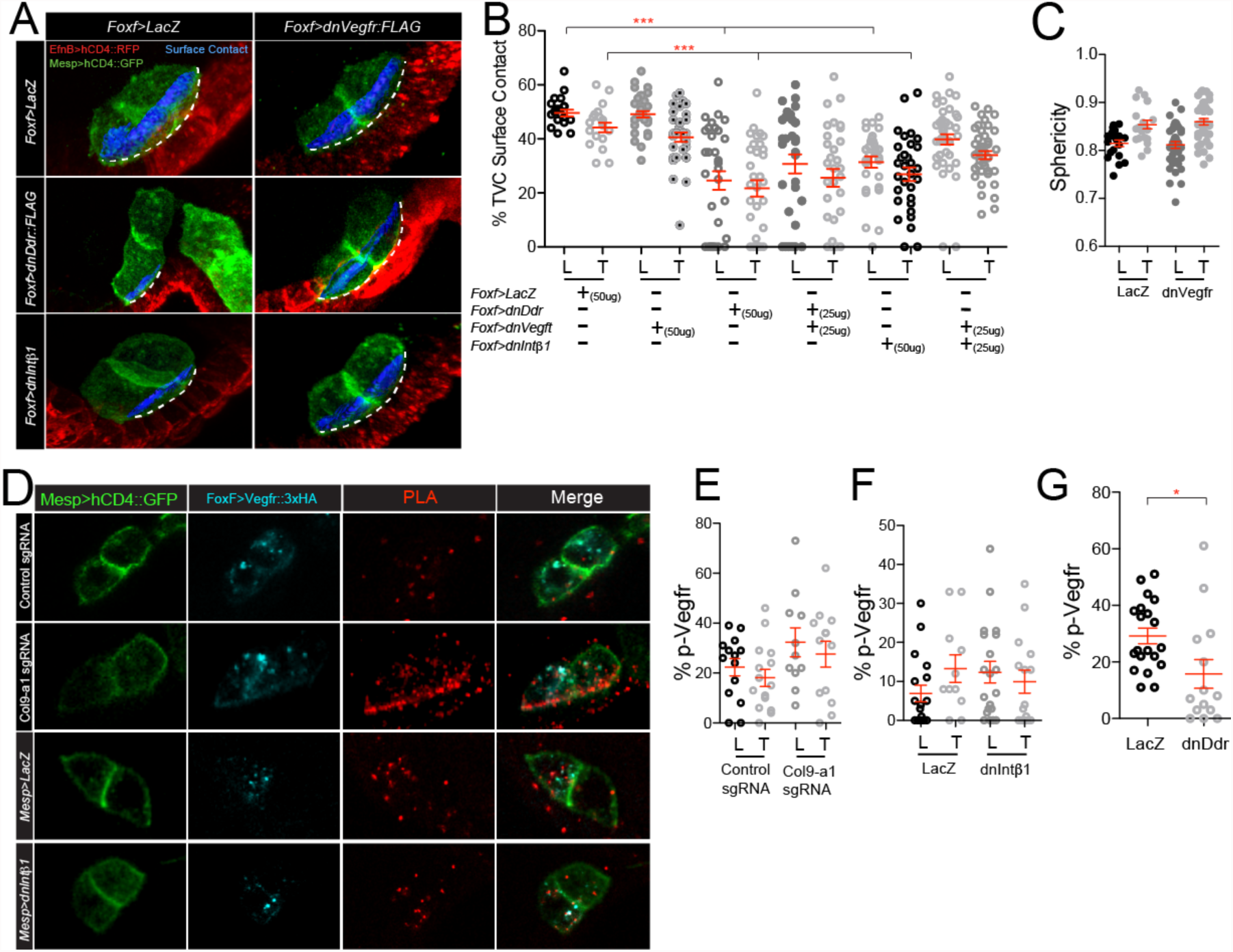
Vegfr modulates TVC adhesion by antagonizing Col9-a1/Ddr/Intβ1. **A**. TVC adhesion under double dn conditions. TVCs are visualized using Mesp>hCD4::GFP, epidermis is visualized using *EfnB>hCD4::tagRFP*. Extent of epidermal surface content is shown with dotted line. Blue surfaces represent portion of TVC in contact with epidermis. **B**. Quantitation of TVC L/T surface in contact with epidermis under conditions that combine dnVegfr and dnDdr or dnIntgB3. Note that this data set originally included the mCherry, dnDdr, and dnIntβ1 conditions that are shown in Figure 2. **C**. Sphericity of leader/trailer TVCs under control and dnVegfr conditions. **D**. PLA assay to detect levels of phosphorylated Vegfr in Col9a-1 CRISPR, dnIntβ1, and dnDdr conditions. **E-F**. Percent phosphorylated Vegfr under perturbation conditions. All images are oriented with leader TVC to the left. L=leader, T=Trailer. * = p<0.05, ** = p<0.005, *** = p<0.0005.

We next tested whether components of the Col9-a1/Ddr/Intβ1-mediated cell-matrix adhesion system modulate Vegfr activity. We performed PLA assays to quantify Vegfr phosphorylation states following CRISPR/Cas9-mediated mutagenesis of *Col9-a1* in endoderm progenitors, and misexpression of either dnDdr, or dnIntβ1 in the TVCs. In control migrating TVCs, we observed most Vegfr::HA signal in cytoplasmic punctae, where phosphorylated Vegfr represented about 20% of the total pool (Figure 5D,E). Both Col9-a1 inhibition and dnIntβ1 misexpression resulted in an increased of Vegfr activity to ~30% of the total pool, with a preferential localization at the ventral membrane contacting the epidermal matrix (Figure 5). On the other hand, dnDdr misexpression decreased Vegfr activity (Figure 5G). Taken together, the results obtained with Col9a1 and Intβ1 are consistent with the hypothesis that cell-matrix adhesion antagonizes Vegfr signaling, thus contributing to a mutual antagonism, possibly modulating cell-matrix adhesion in migrating TVCs. By contrast, the unexpected effects of dnDdr suggest that Ddr is also required independently of its effects on cell-matrix adhesion to promote Vegfr signaling.

### Asymmetric exposure to Col9-a1, and Ddr signaling establish and maintain collective TVC polarity

Previous studies showed that asymmetric contacts between the TVCs and surrounding tissues contribute to canalizing TVC behavior towards collective leader/trailer polarity and directed migration^13^. Specifically, the prospective leader cell, born more ventrally, initially contacts the endoderm, whereas the future trailer interacts with the mesenchyme. This asymmetry opens the possibility that an endodermal product, such as Col9-a1, contributes to establishing collective leader-trailer polarity prior to migration. To determine if differential contact with the endoderm determines asymmetrical exposure to Col9-a1, we imaged Col9-a1::GFP deposition in the ECM and quantified colocalization with the TVC membranes at early stages, prior to and shortly after the onset of TVC migration (Figure 6A). *Nkx2-1*-driven Col9-a1::GFP gradually accumulated in the ECM between stage 17 and 19, and clearly labeled the TVC/epidermis interface by stage 21, when TVCs have begun their migration. The prospective leader cell membrane initially colocalized more extensively with Col9-a1::GFP foci, compared to the presumptive trailer (Figure 6A). This suggests that Col9a1::GFP does not diffuse extensively, and potentially provides a polarizing signal to the TVCs through asymmetric exposure. As cells initiated migration, endoderm-derived Col9-a1::GFP colocalized similarly with leader and trailer. This implies that Col9-a1 induced polarity must be established prior to the onset of migration, and subsequently maintained through migration.

**Figure 6.**
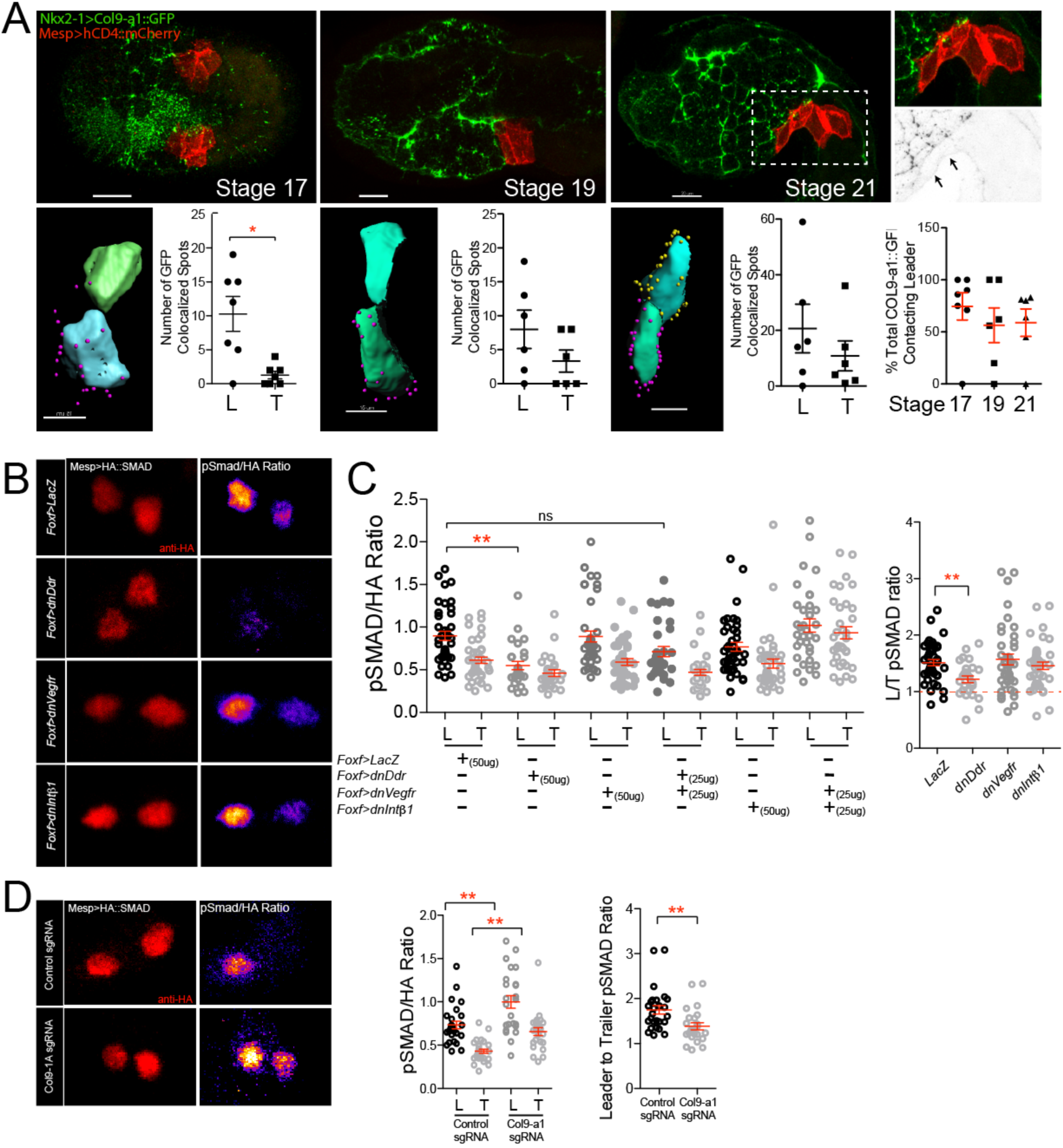
Asymmetric exposure to Col9-a1 polarizes TVCs through an adhesion-independent function of Ddr. **A**. Distribution of Col9-a1::GFP fusion expressed in the endoderm at stages 17-21, prior to and at the onset of TVC migration, respectively. Boxed and expanded region shows localization of Col9-a1::GFP fusion ventral to the migrating TVCs. Calculation of total exposure of leader or trailer to endodermal collagen at each embryonic stage. Presumptive leader is shown in blue, presumptive trailer in green. Purple spheres indicate collagen spots proximal to the leader TVC, yellow spheres are proximal to the trailer TVC. Graphs show total number of collagen contact sites for leader or trailer at each given time point. Total number of Col9-a1::GFP spots contacting leader TVC over time. **B**. Quantitation of pSmad activity under conditions perturbing TVC adhesion. Ratios between the pSmad and HA channels are calculated by dividing the pSmad channel by the HA channel. Ratio is color-coded using the Fire Look Up Table (LUT) with lighter colors indicating a higher ratio. **C**. Normalized levels of pSmad in L/T under perturbation conditions and ratios of total leader:trailer pSmad levels. **D**. pSMAD staining of control migrating TVCs and TVCs under endodermal Col9-a1 CRISPR mutagenesis conditions. L:T ratios and normalized L/T pSmad levels are shown to the right. **E**. All images are oriented with leader TVC to the left. L=leader, T=Trailer. * = p<0.05, ** = p<0.005, *** = p<0.0005.

Collective TVC polarity is evidenced by morphological differences between the leader and trailer cells^13, 16^, but no clear molecular marker has been identified. We previously proposed that TVCs are exposed to varying levels of BMP-Smad signaling^48^. To test whether BMP-Smad signaling is leader-trailer polarized in *Ciona* TVCs, we developed a biosensor by humanizing the C-terminus of *Ciona robusta* Smad1/5/8 and adding an HA tag at the N-terminus. Using both an anti-HA and an anti-phospho-SMAD5 antibody, which cross-reacts poorly with the endogenous *Ciona* protein, we quantified BMP-Smad signaling in the B7.5 lineage as the pSmad/HA ratio, following expression of this HA::Smad1/5/8^Hs^ sensor using the *Mesp* enhancer. This sensor responded to defined perturbations of the BMP-Smad pathway (Figure S6), indicating that it can serve as a reliable readout for endogenous signaling. In control embryos, the PSmad/HA levels were ~1.75 times higher in the leader compared to the trailer cell (Figure 6B,D). Therefore, we used changes in pSmad/HA levels to assay the roles of Col9-1, Ddr, Intβ1 and Vegfr signaling in establishing and maintaining collective leader/trailer polarity.

We first quantified changes in BMP-Smad signaling following perturbations of Ddr, Intβ1 and Vegfr functions. TVC-specific expression of dnDdr reduced the absolute PSmad/HA ratios, and abolished differences between the leader and trailer cells, suggesting that Ddr promotes BMP-Smad signaling and contributes to establishing and maintaining TVC polarity. By contrast, misexpression of neither dnIntβ1 nor dnVegfr did alter pSmad/HA levels or asymmetry in the TVCs (Figure 6B,C). In an attempt to reveal cryptic activity, and because Vegfr appeared to antagonize the functions of Ddr and Intβ1 in cell-matrix adhesion, we tested whether dnVegfr or dnIntegB3 could modulate the effects of dnDdr on BMP-Smad signaling. In neither case did we observe significant differences compared to dnDdr alone. Taken together, these results suggest that Intβ1 and Vegfr act primarily upon cell-matrix adhesion, whereas Ddr coordinates collective polarization and adhesion, by independently regulating each pathway.

Finally, since endoderm-derived Col9-a1 is necessary to activate Ddr in the TVCs, and newborn TVCs may be differentially exposed to extracellular Col9-a1, we probed BMP-Smad signaling following CRISPR/Cas9-induced *Col9-a1* mutagenesis. Unexpectedly, *Col9-a1* inactivation in the endoderm caused a general increase of pSmad/HA levels in both leader and trailer cells, and a slight reduction in leader-trailer ratio (Figure 6D). These observations are seemingly inconsistent with a role for Col9-a1-activated Ddr in promoting BMP-Smad signaling in a polarized fashion. However, because Col9-a1 acts in the extracellular milieu, we hypothesize that Col9-a1 may contribute to the sequestration of BMP ligands, as shown for type IV collagens in *Drosophila^49^*, thus possibly reducing the availability of ligands for signaling via BMP receptors at the TVC membrane. Although these hypotheses remain to be tested, our results are consistent with a model whereby Col9-a1 regulates BMP-signaling via an incoherent feedforward signaling circuit, which consists of an inhibition via ligand sequestration that is partially compensated by activation via Ddr (Figure 7).

**Figure 7.**
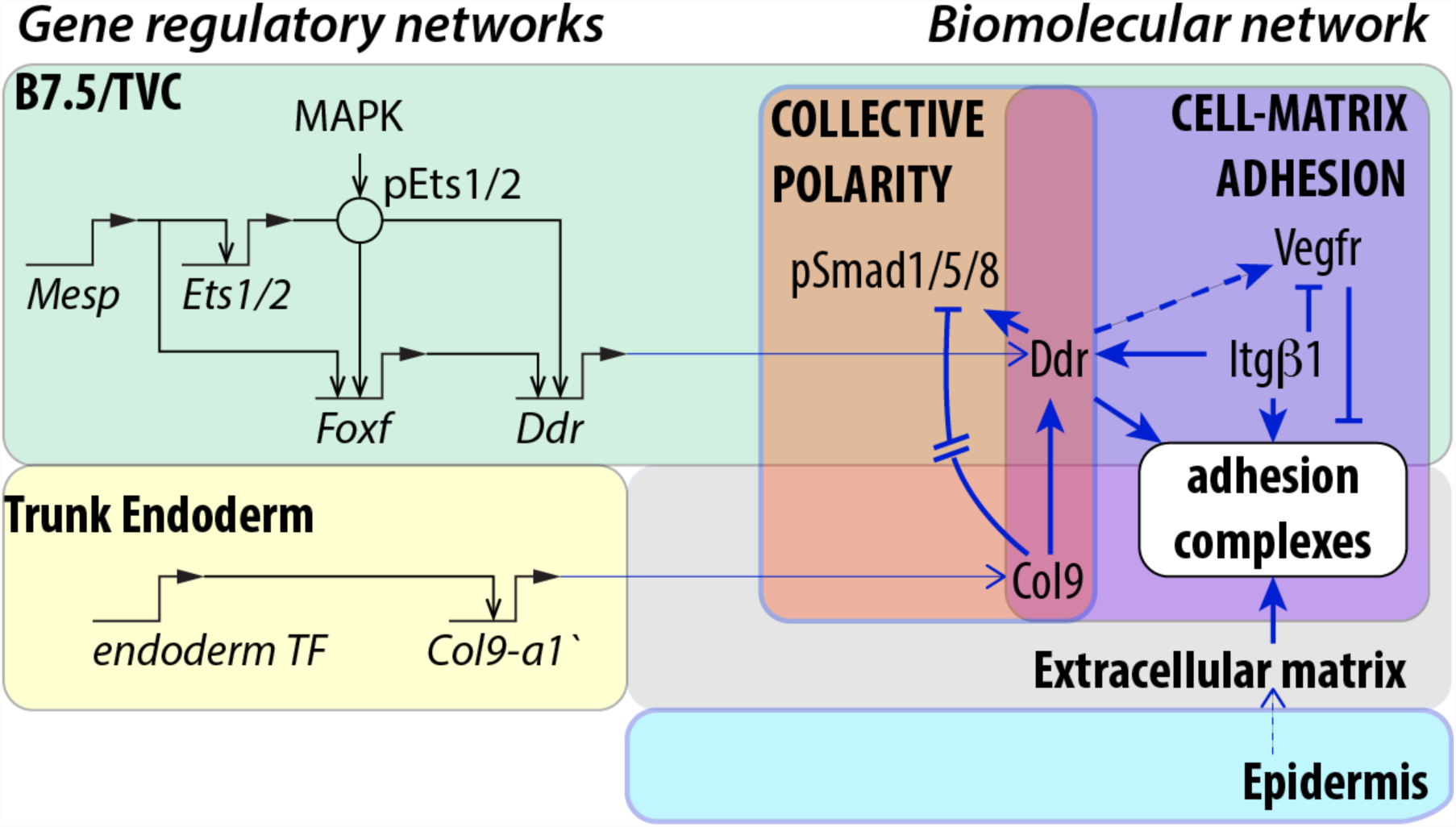
Integration of transcriptional and biomolecular networks to produce directional migration. Intersection of fate-specification through the cardiac GRN in the B7.5/TVC lineage (green) and downstream cellular effectors regulating ECM adhesion and polarity (purple and orange) produces directional migration. Black arrows show transcriptional regulation, blue arrows show activity regulation. Gene regulatory connections are after^14-16, 18, 21^. Exposure to extracellular Col9-a1 activates Ddr and Intβ1 while blocking Vegfr activity. Solid blue lines show interactions confirmed in this report. Dashed lines show indirect relationships.

## DISCUSSION

In embryos, the behaviors of progenitors cells emerge from the integration and coordination of inputs from intrinsic factors, including transcriptional regulation of cellular effector genes, and biochemical and mechanical extrinsic determinants1. Here, we showed that, in migratory cardiopharyngeal progenitors of the tunicate *Ciona*, the transcription factor Foxf is necessary to upregulate the collagen receptor Ddr, which senses the extracellular matrix and functions to promote both cell-matrix adhesion and collective polarity through presumably independent pathways.

This study corroborates preceding transcriptional profiling results showing that Foxf, an early component of the cardiopharyngeal regulatory network, which was previously shown to impact protrusive activity by regulating the expression of the small GTPase Rhod/f^16^, also connects the GRN to cellular effectors capable of sensing and responding to the extracellular milieu. This suggests that Foxf regulates a cell-matrix adhesion module, as previously proposed^16^. Notably, FoxF1 was shown to regulate Integrin-β3 expression and cell-matrix adhesion in the mouse^50^, suggesting an ancient and conserved connection between Foxf homologs and the regulation of cell-matrix adhesion.

Previous studies used a dominant negative form of Foxf, with the WRPW repressor motif fused to the DNA-binding domain, to propose that Foxf activity controls TVC migration^15^. This reagent produced more severe migration phenotypes than our newly developed CRISPR/Cas9 assay, which nevertheless strongly inhibits *Foxf* expression (Gandhi et al., 2017^36^ and data not shown). This discrepancy suggests that the repressor form of Foxf caused non-specific defects, such as inhibiting the targets of other Forkhead box family factors expressed in the B7.5 lineage^16^, which may not be affected by CRISPR-induced Foxf loss-of-function. Future studies will clarify the functions of these other Fox factors at the transcription-migration interface, and their possible interactions with Foxf.

While TVC-specific transcriptional inputs control the expression of cell-matrix adhesion determinants, asymmetric adhesion to the extracellular matrix was previously shown to polarize the founder cells and permit localized TVC induction, and thus *Foxf* expression^*51*^. Precocious misexpression of dnDdr using the *Mesp* driver interfered with this integrin-dependent TVC induction, thus supporting the notion that dnDdr interferes with cell-matrix adhesion (Figure S7). This illustrates the notion that cell fate specification and subcellular processes are dynamical and interdependent components of the transcription-cell behavior interface, which is best understood as an integrated system.

Our results indicate that at least two putative collagen receptors, Ddr and Intβ1, contribute to cell-matrix adhesion during TVC migration. Discoidin-domain receptors and integrins have been showed to interact during cell adhesion to collagen matrices^34, 52, 53^. Whereas the function of integrins as cell-matrix adhesion receptors mechanically coupled to the cytoskeleton is well described^54, 55^, Ddr homologs bind collagens independently of integrins^38, 53, 56^, and they appear to function as weak adhesion molecule and/or an ECM sensor, which can promote cell-matrix adhesion through other pathways, including integrins^34, 57^.

A prominent feature of our model is the functional antagonism between Vegfr signaling and Ddr/Intβ1-dependent cell-matrix adhesion. Although ligands and signal transduction pathways remain to be characterized, the antagonism appears to occur in part through local inhibition of Vegfr at the ventral membrane, where Ddr and Intβ1 bind the ECM (Fig. 5). While functional interactions between VEGF receptors and integrin complexes are largely context-dependent^45^, VEGFR2 and αvβ3 integrins were found to directly interact through their cytoplasmic tails^58^ and integrin activity can promote VEGFR2 degradation^46, 47^. Conversely, VEGFR2 activity has been linked to turnover of integrin-based focal adhesion^59^. It is thus possible that Vegfr can induce endocytosis of integrin complexes, thus weakening cells’ adhesion to the basal lamina. Remarkably, in Ciona cardiopharyngeal progenitors, the Ddr/Intβ1-Vegfr antagonism contributes to positioning cells between the endoderm and the epidermis, a mesodermal hallmark in triploblastic animals.

Besides their role in cell-matrix adhesion, the ECM-sensing properties of Ddr contribute to the establishment of distinct leader and trailer states, as a result of differential exposure of to extracellular collagen. This is reminiscent of the roles of ECM components in establishing of leader cell states in wound scratch assays *in vitro*^60^, and in guiding endomesoderm migration during gastrulation^61^. Moreover, DDR1 can interact with the PAR complex to polarize the actin cytoskeleton in individual cells^62^. Future studies will uncover the molecular pathways by which Ddr control collective polarity, and the elusive cell-cell communication mechanisms that we must invoke, by analogy with other systems^63, 64^, to explain maintenance of leader-trailer polarity throughout migration.

We reported differential BMP-Smad signaling as a reliable read-out of polarized leader/trailer states. Ddr and Col9-a1 are required for this anisotropic BMP signal, which appeared largely independent of Intβ1-mediated cell-matrix adhesion and Vegfr signaling. This indicates that Ddr acts as a dual function receptor, promoting cell-adhesion and transducing polarity information through independent pathways. Although Ddr is activated by exposure to Col9-a1 and promotes BMP-Smad signaling, loss of Col9-a1 function increased BMP-Smad signaling. To reconcile these seemingly inconsistent results, we hypothesize that BMP ligands are sequestered by extracellular collagen and inaccessible for signaling, as reported in the Drosophila ovary^49^. In this model, extracellular Col9-a1 would thus control BMP-Smad signaling through an incoherent feedforward signaling circuit, whereby activation of Ddr-mediated signaling would compensate ligand sequestration in the extracellular matrix. Future studies will test these possibilities and explore the signal transduction pathways connecting Ddr to BMP-Smad signaling.

## Acknowledgements

We thank Justin Le Lorier for cloning the HA::Smad1/5/8^Hs^ sensor. This work was supported by NIH F32 GM108369-01A1 post-doctoral fellowship to Y.B., NIH F32 GM105216-01A1 post-doctoral fellowship to S.G., and NIH/NIGMS R01 GM096032 award to L.C.

## Authors’ contributions

Y.B. and L.C. designed the experiments, Y.B., S.G., S.B., and W.W. performed the experiments. Y.B. and L.C. wrote the paper.

## Materials and Methods

### Electroporation and transgene expression

*Ciona robusta* (formerly known as *Ciona intestinalis* type A^65^) adults were purchased from M-Rep, San Diego, Ca. Gamete isolation, fertilization, dechorionation, and electroporation were performed as described^66-68^. The amount of DNA electroporated varied from 10μg to 80μg. Animals were reared at 16 to 20°C. For proximity ligation assays, embryos were fixed in cold 100% methanol for 10 minutes. For *in situ* hybridization experiments, embryos were fixed for 2 hours in 4% MEM-PFA, dehydrated in an ethanol series and stored in 75% ethanol at −20°C as described^66, 69^. Embryos used for direct visualization of fusions were fixed in 4% MEM-FA for 30 minutes, cleared with an NH4Cl solution, and imaged using a Leica SP8 X Confocal microscope.

### Molecular Cloning

Coding sequences of Ddr (KH.C9.371) Vegfr (KH.C14.345), Fgfr (KH.S742.2), and (Egfr(KH.L22.45) were amplified from mid-tailbud *Ciona* cDNA libraries. To generate dominant negative constructs, cytoplasmic kinase domains were removed by truncating the sequence after the transmembrane and extracellular domains. Primer lists are given in Supplementary Table 1. To subclone larger protein-coding fragments we used a multi-fragment assembly approach using the NEB InFusion homologous recombination kits. Coding regions that were greater than 2kb in length were subdivided into overlapping kernels and ligations of up to 4 kernels were performed.

### Live imaging and TVC tracking

To generate 4D datasets, 4.5 hpf old embryos were mounted on glass bottom microwell petri dishes (MatTek, part# P35G-1.5-20-C) in artificial seawater. Plates were sealed by piping a border of a mix of Vaseline and 5% mineral oil by volume (Sigma, item #M841- 100 ml) and covering with a 22x22 Fisherbrand Cover Glass (item # 12-541-B). Embryos were imaged on a Leica inverted SP8 X Confocal microscope every 3.5 minutes for 4 to 5 hours. B7.5 lineage nuclei and epidermal cell membranes were visualized using Mesp>H2B::GFP and EfnB>hCD4::mCherry^13^, respectively, and TVC migration was tracked using Imaris Software.

To express TVC position during migration in 3D we subdivide the embryo into quadrants using two conceptual orthogonal planes that bisect the developing embryo. The position of the midsagittal plane is determined by the midline of the epidermis visible using the EfnB>hCD4::mCherry transgene. The frontal plane is orthogonal to the mid-sagittal plane and passes through the nucleus of the anterior ATM and the future position of the palps, the most anterior point of the embryo. Imaris Bitplane is used to calculate a vector line based on position of GFP+ TVC nuclei.

### Proximity Ligation Assay

To visualize phosphorylated RTKs each receptor was subcloned into an expression vector driven by either the *Mesp* (early expression) or *Foxf(TVC)bpFog* (late expression) enhancer and tagged with 3x hemagglutinin (HA). Expression vectors were electroporated into fertilized *Ciona robusta* embryos, which were raised at room temperature to desired stages as indicated. Embryos were fixed by washing in 1ml of cold 100% methanol for 10 minutes and stored in 100% methanol −20 C until processed. Fisherbrand slides were coated with 0.1% w/v Poly-L-Lysine (Sigma P8920) two times and allowed to dry. Embryos were rehydrated in a 0.1% PBT (0.1% Tween-20 in PBS): MeOH series of 3:7, 1:1, 7:3, 100% PBT for 15 minutes and mounted on the Poly-L-Lysine coated slides. Slides were blocked in three drops of Duolink Blocking reagent for 30 minutes at 37°C. Rabbit Anti-HA and mouse anti-pTyr were used to visualize phosphorylated RTKs and a rat anti-HA antibody was used to visualize total amount of RTK expressed in the embryo. Antibodies were used at a concentration of 1:500 diluted in Duolink Antibody Diluent. Slides were incubated with primary antibody for 1 hour at 37 degrees and washed 2x in home made Duolink Buffer A for 15 minutes each in a Coplin Jar. Secondary Duolink anti-mouse and anti-rabbit antibodies conjugated to a DNA probe were used as directed by the Duolink manual. AlexaFluor-555 anti-rat secondary antibody (Life Technology #A21434) was added at a 1:1000 dilution to the mix of secondary antibodies to detect total amounts of HA. 100μl total of secondary antibodies were used per slide. Slides were incubated for 1 hour at 37°C, washed two times in home-made Duolink Buffer A for 15 minutes each in Coplin jars. Ligation reactions were carried out as recommended in the Duolink manual, but extended to 1 hour at 37°C. Slides were washed again two times in Duolink Buffer A for 15 minutes each in Coplin jars. Polymerase reaction was carried out as recommended, and extended to 2 hours at 37°C. Slides were washed two times in home-made Duolink Buffer B for 15 minutes each and one time in 0.01% Buffer B for 15 minutes. 15 μl Duolink Mounting Media was added to each slide, 22x22 Fisherbrand coverslips were placed on top and the slides were sealed with clear nail polish.

### Fluorescent in situ hybridization – FISH-IHC

RNA hybridization probes were generated from late tailbud derived cDNA by subcloning the coding regions of the genes of interest into the pCRII-TOPO dual-promoter cloning vector (Invitrogen, KH461020) and PCR was performed to confirm the orientation of the insert. Coding regions were amplified using M13 Forward and Reverse primers and the PCR product was used as a template to synthesize RNA probes labeled with either digoxigenin or fluorescein using the SP6 polymerase. Primers for probe generation are listed in Supplemental Table 1. Single and double FISH were performed as described^66, 69^.

### CRISPR/Cas9 - guide RNA design

Single guide RNAs (sgRNAs) targeting the *Col9-a1* locus (KH.C8.248) were designed and tested essentially as described^36, 43^. To test the efficiency of the sgRNAs, we first mutated the Col9-a1 locus using the ubiquitous *Ef1a>nls::Cas9::nls* and the *U6>sgRNA* constructs. Cutting efficiency was calculated based on Sanger sequencing of the sgRNA target region as described^36^. Cutting efficiency of *Col9-a1* sgRNAs peaked at 0.8 and 1.1 for sgRNAs targeting the 1^st^ and 29^th^ exon, respectively. We targeted the *Col9-a1* locus by combining two high efficiency Col9-a1 sgRNAs, expected to generate large deletions. To target the *Col9-a1* locus in endoderm progenitors, we used the vegetal hemisphere enhancer from *FoxD* to drive nlns::Cas9::nls expression. 25μg of Cas9-expressing vector and a total of 80μg of sgRNA-expressing constructs were used for each experiment. As a control, sgRNAs targeting the late-expressed *Ebf* gene were used throughout the manuscript.

### Design and cloning of short hairpin RNAs (shRNAs)

shRNA were designed as described^22, 69^. We designed the Ciona short hairpin microRNA (Ci-shmiR) cassette based on the primiR structure of Cirobu.mir-2213-a^70^. The Ci-shmiR cassette (aaa gcggccgc aaa gctagca taa tga acttcgtggccgtcgatcgtttaaagggaggtagtga ggtacc tctagt ggatcc [cgcggcgctaggttcgtttaatggtctaaaaatcaGagcgtttagt**GTTTG**<u>gagaccg</u>agagag<u>ggtctc</u>actaaaactgcgcttattatcttctacgaacctgtaagtggc] agatct ggccgca ctcgag tttg atgaattccagctgagcg) was cloned downstream of the Mesp enhancer. Hairpins were cloned into the Ci-shmiR cassette using BsaI. Cloned hairpins were validated by testing for a knockdown of target genes tagged with a GFP compared to cell markers that are not fused to gene targets. Hairpins that knocked down the reporter were further validated by *in situ* hybridization for the target gene.

### BMP-Smad biosensor, pSmad staining and quantification

The Mesp>HA::Smad1/5/8^Hs^ was built by replacing the C-terminus of *Ciona* Smad1/5/8 by that of human SMAD5, using standard cloning procedures. A monoclonal Rat anti-HA antibody was used to evaluate total levels of expressed protein and to normalize pSMAD levels across multiple embryos. Polyclonal Rabbit anti-phospho-SMAD5 was used to label phosphorylated HA::Smad1/5/8^Hs^ proteins, and anti-beta-galactosidase against the protein product of Mesp>LacZ was used to visualize the B7.5 lineage. To quantitate pSMAD levels as ratios, we use the anti-beta-galactosidase signal to identified TVC nuclei as spots in Bitplane Imaris. We then took the quantified the average fluorescence of anti-pSmad and anti-HA in the leader and trailer TVC. Ratios of pSmad to HA were calculated to normalize for variable transgene expression. The sensor’s response to BMP-Smad signaling was validated using expression of published constitutively active BMP receptor and Noggin^48^, an extracellular inhibitor (Figure S6).

### Image acquisition

All images were acquired using the Leica SP8 X WLL Confocal microscope using the 63x oil immersion lens. Z-stacks of fixed embryos were acquired at the system optimized Z-step, 512x512 resolution, 600 Hz, and bi-directional scanning. Multiple HyD detectors were used to capture images at various wavelengths.

### Morphometrics analysis and surface contact calculation

The membrane marker Mesp>hCD4::GFP was used to segment the TVCs and derive morphometric measurements such as sphericity, area, and volume in Bitplane Imaris using the Cell function. Z-steps were normalized to achieve equal voxel size in X, Y, and Z planes. TVCs were then segmented and resulting cells were exported to separate surfaces. Another surface was created for the epidermis using the EfnB>hCD4::tagRFP or mCherry marker. Distance transformation was performed on each TVC and the epidermal surface was then used to mask the distance from the epidermal surface to the distance-transformed surface of each TVC. A new surface representing the surface of contact between the TVC and the epidermis was created using the masked distances. To calculate the percent of the TVC surface in contact with the epidermis the area of the surface contact was divided by two and then divided by the total area of the TVC.

### Colocalization

To identify spots of colocalization the CoLoc module of Bitplane Imaris was used to create a colocalization channel. The threshold of each image was set by the signal of interest, e.g. to find colocalization between the Mesp>hCD4::mCherry and Nkx2-1>Col9-a1::GFP the threshold was set to include the red channel. A separate channel was then created based on the colocalization and each area of that channel was transformed into a spot. We then calculated the number of spots close to the leader and trailer by identifying spots within 5 microns of the TVC surface.

---

**Supplemental Figure 1.**
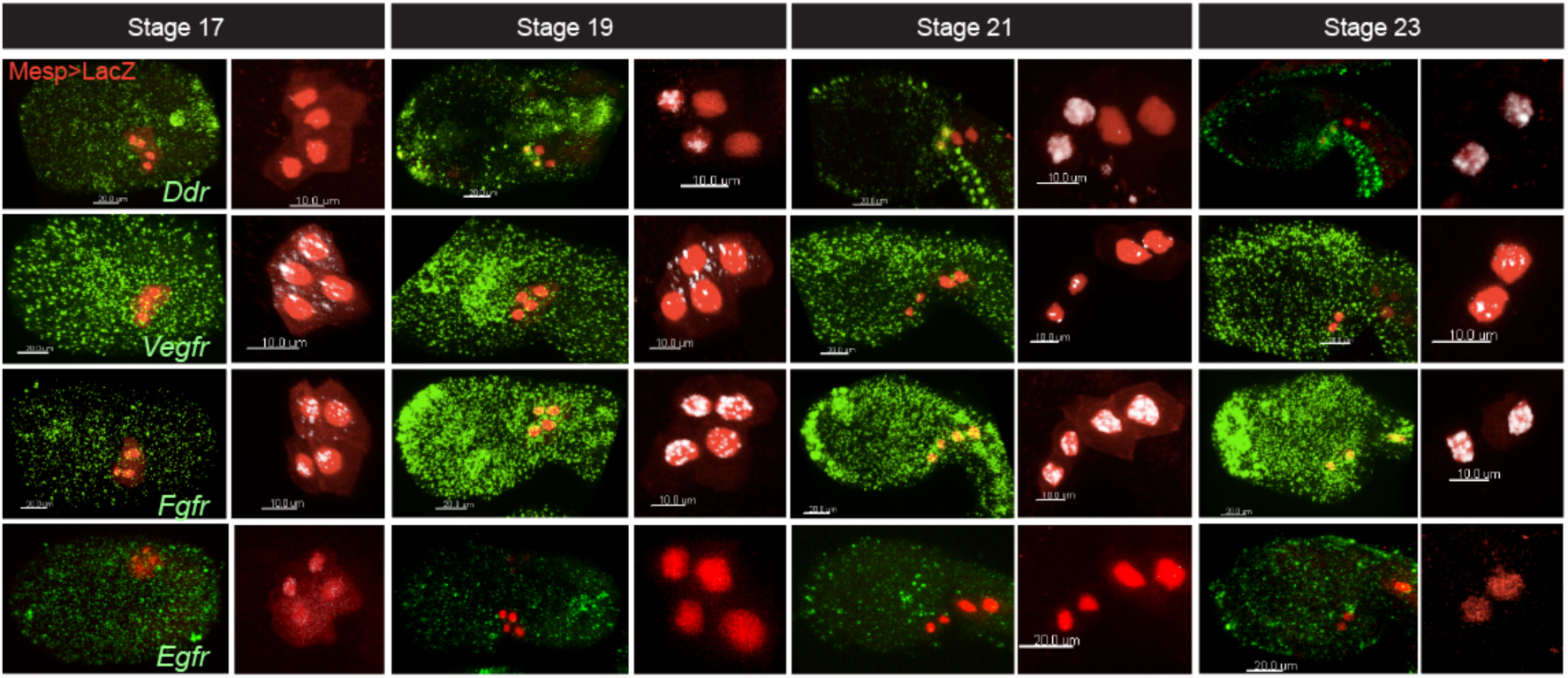
RTK expression during TVC migration. Fluorescent *in situ* hybridization for *Ddr, Vegfr, Fgfr,* and *Egfr* at indicated developmental stages. B7.5 lineage is marked with *Mesp>LacZ* and stained for beta-galactosidase (red in the micrographs). Close ups show colocalization of transcripts with B7.5 lineage nuclei.

**Supplemental Figure 2.**
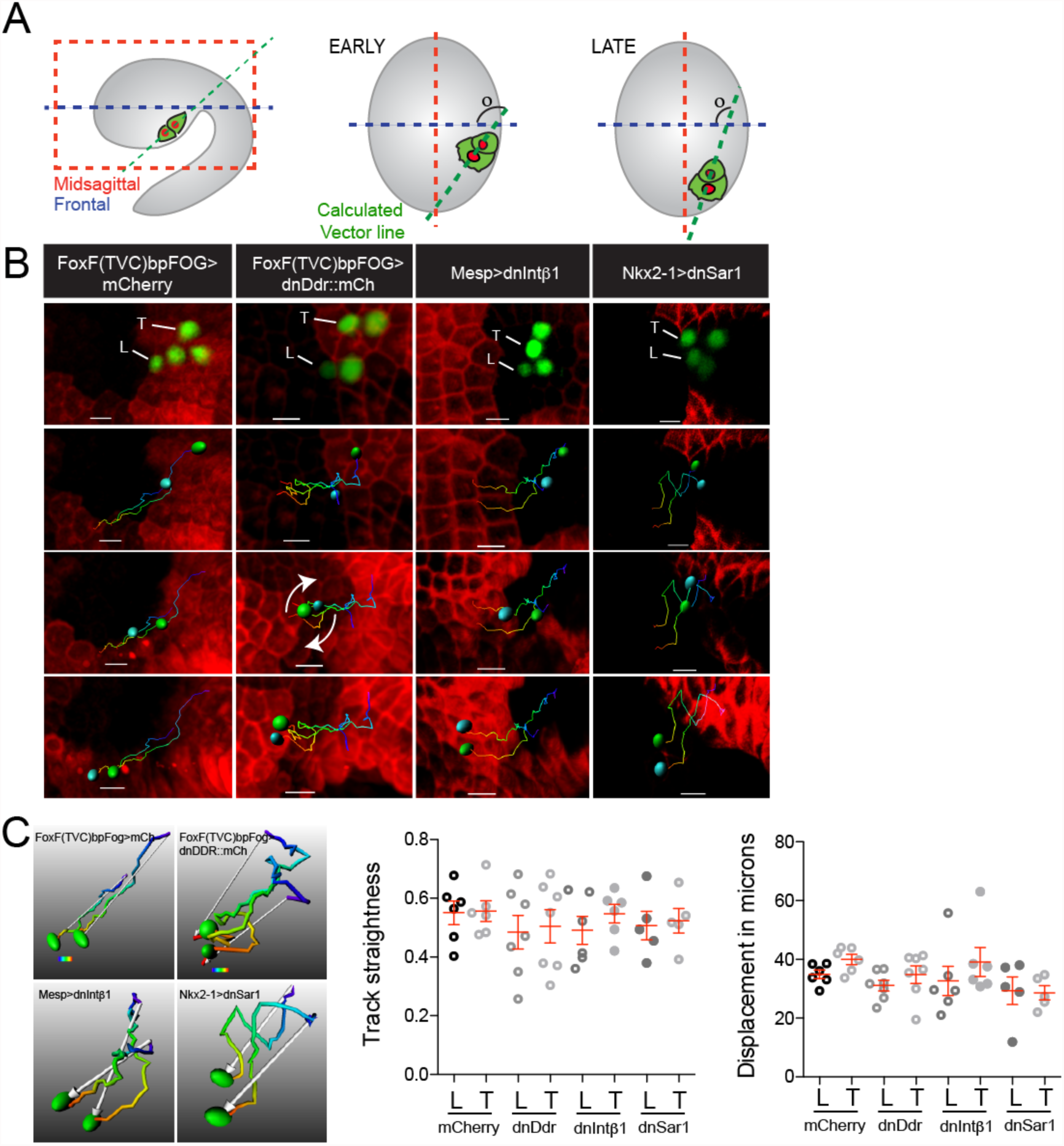
Quantitative analysis of TVC migration. **A.** Schematic of calculating TVC position relative to frontal (blue) or sagittal (red) planes. The vector line (green) is calculated using the axes of the TVC nuclei. **B.** Positional tracking of TVC migration under perturbations of ECM adhesion. Graphs show average positions of TVCs relative to orthogonal planes and standard deviation at each time point. **C.** Rendered images of total displacement of the TVC under adhesion perturbation. White arrows point to the final position of TVCs prior to the first asymmetric division. TVC tracks are time code blue for early time points and red for late time points. **D.** Average track straightness and total displacement of TVCs.

**Supplemental Figure 3.**
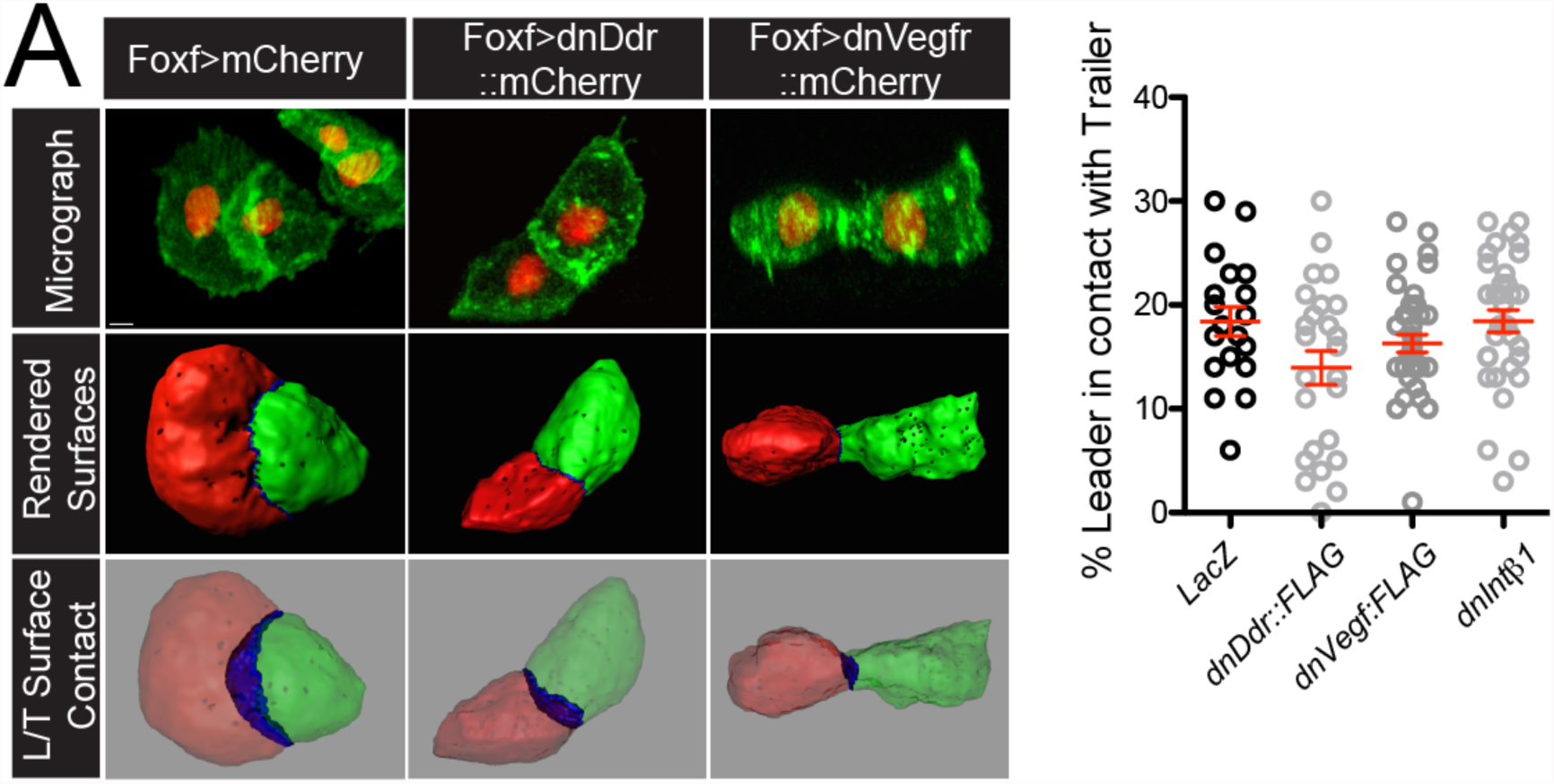
Ddr is required for leader/trailer cell-cell junction. Micrographs of TVCs undergoing migration (top row). Nuclei are marked with *Mesp>H2B::mCherry,* cells are visualized using *Mesp>hCD4::GFR* TVC pairs are segmented using Imaris Bltplane software, middle row. Leaders are in red, trailers in green. Surface of contact between leader and trailer is calculated (bottom row). Average size of the cell-cell junction is reported as percent surface of leader in contact with the trailer.

**Supplemental Figure 4.**
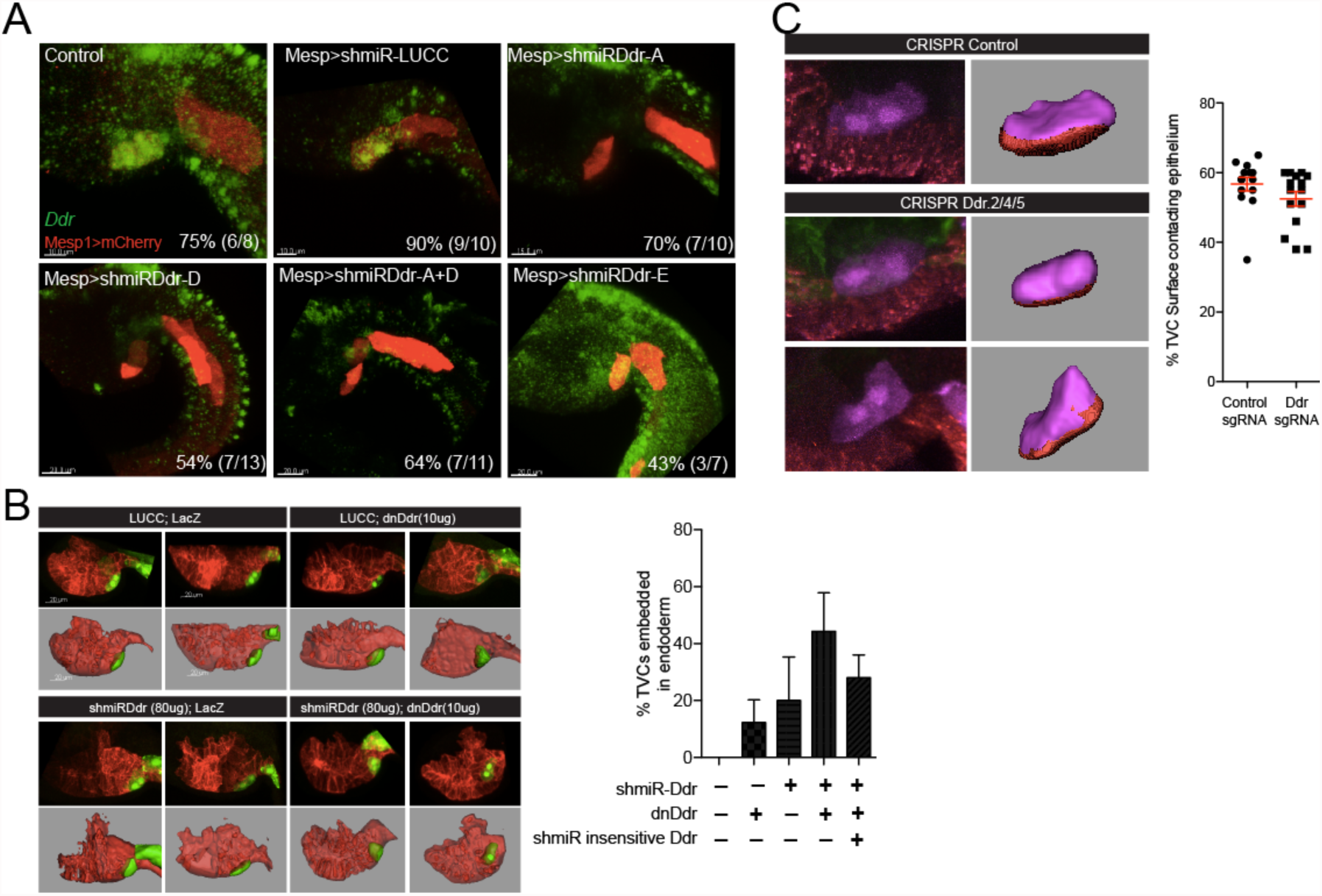
Ddr loss-of-function phenotypes. A short hairpin microRNA (shmiR) knockdown of Ddr transcripts. B7.5 lineage is marked with *Mesp>mCherry.* **B.** Sensitized shmiR test. Two representative embryos are shown for each condition. Top rows show raw data, bottom rows show rendered images. Graph shows % TVC that become embedded in the endoderm as a result of detachment from the epidermis. **C.** Mutagenesis of the *Ddr* locus using CRISPR. *EBF* CRISPR is used as control. Graph shows average percent of the TVC pair surface in contact with the epidermis.

**Supplemental Figure 5.**
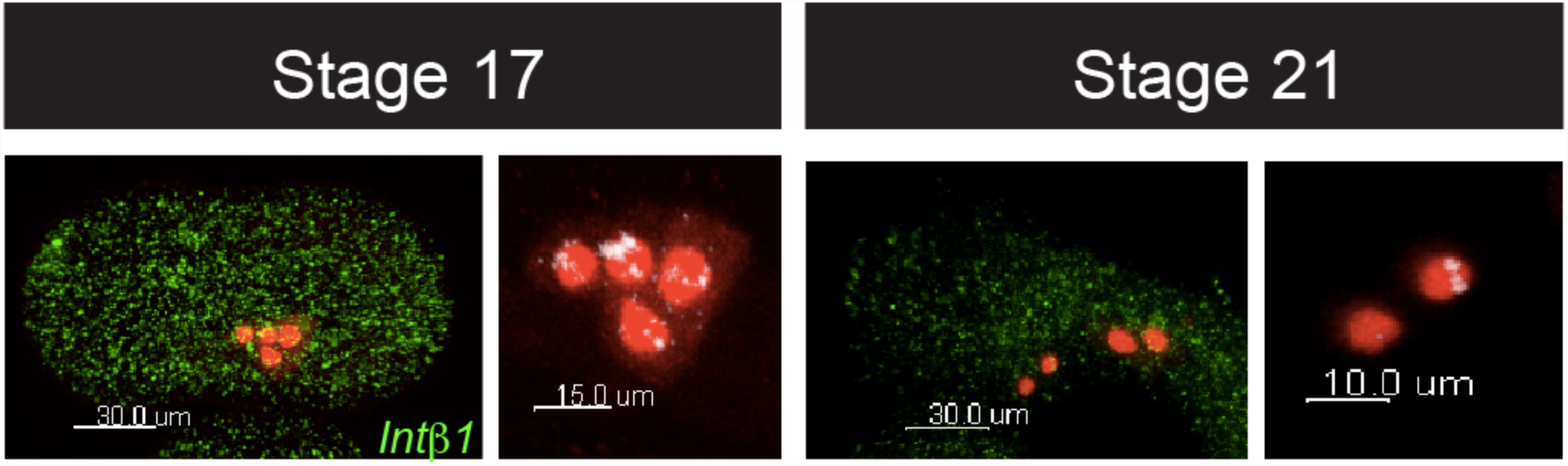
IntgBI expression in the B7.5 lineage. *In situ* hybridization of RNA probe against Intβl transcripts. B7.5 lineage is marked with *Mesp>LacZ.*

**Supplemental Figure 6.**
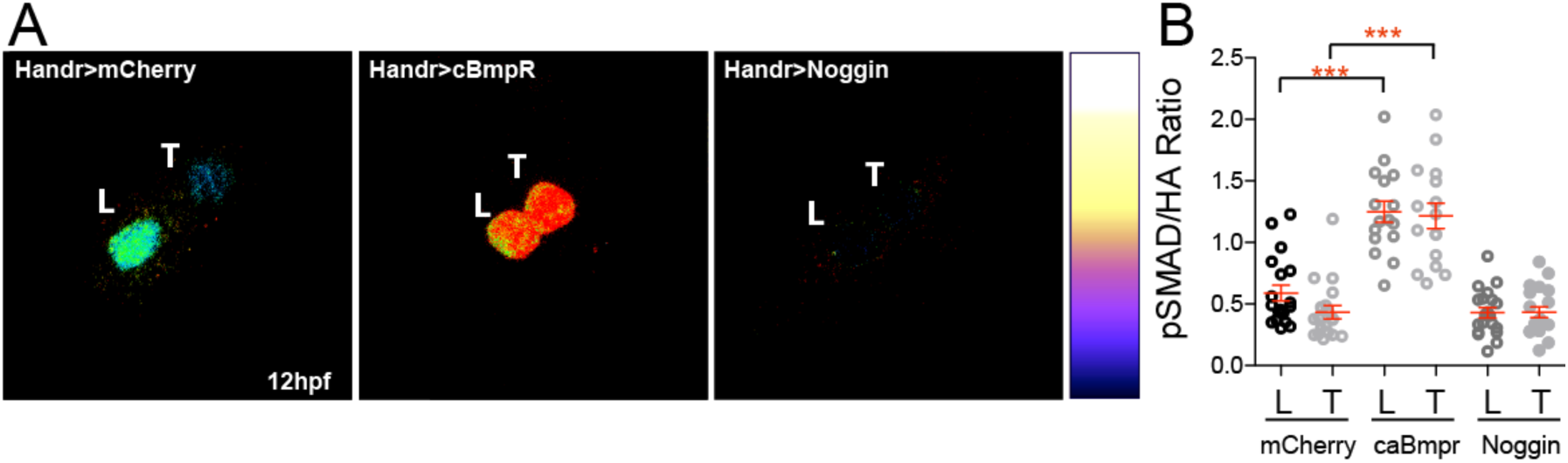
Asymmetric induction of BMP-pSmad activity in migrating TVCs. **A.** Ratios of detected pSmad staining to the total HA levels detected. Levels can be manipulated through addition of a consitituitivley active BMP receptor (Handr>caBmpR) or by expression of the BMP antagonist Nogging (Handr>Noggin). **B.** Calculated ratios of pSmad/HA under conditions altering BMP-pSmad activity levels.

**Supplemental Figure 7.**
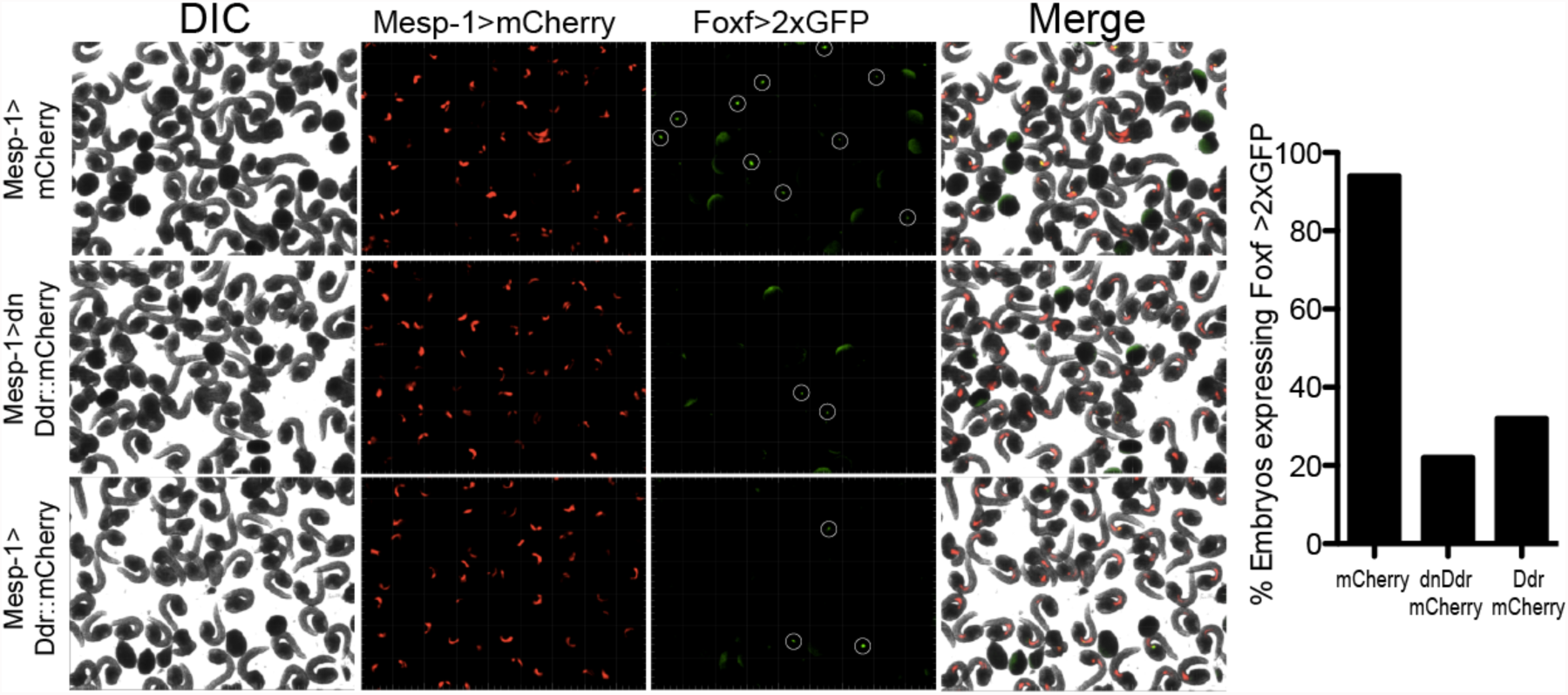
Early expression of Ddr blocks TVC fate induction. Embryos electroporated with Mesp-driven mCherry, a dominant negative Ddr (dnDdr) or full-length Ddr are assayed for expression of Foxf-driven GFP. DIC channel shows embryonic developmental stage. Foxf>2xGFP expressing TVCs are circled in the green channel.

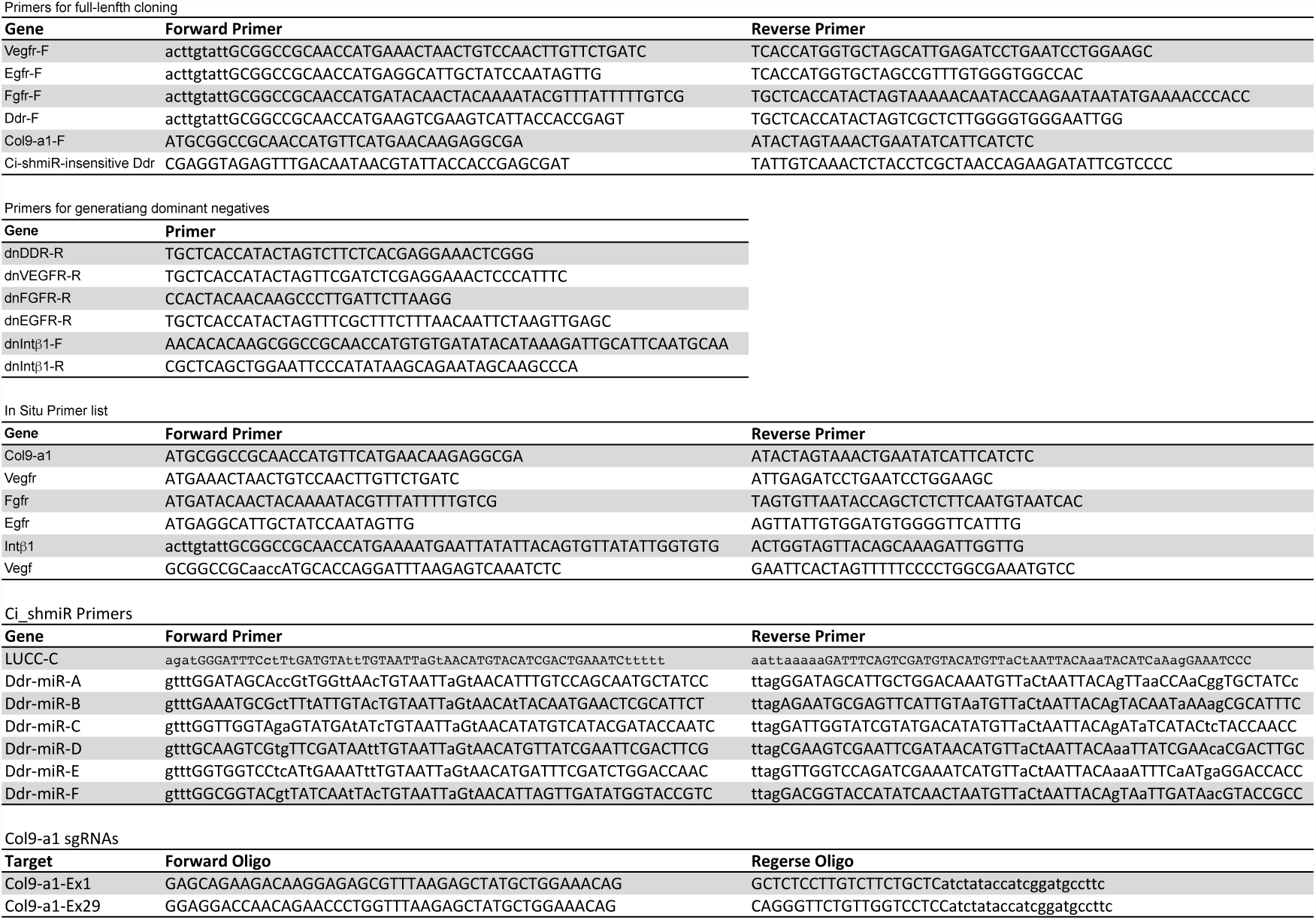
Supplemental Table 1. Primers used for cloning.

